# Real-time visualization of mRNA synthesis during memory formation in live animals

**DOI:** 10.1101/2021.09.01.458476

**Authors:** Byung Hun Lee, Jae Youn Shim, Hyungseok C. Moon, Dong Wook Kim, Jiwon Kim, Jang Soo Yook, Jinhyun Kim, Hye Yoon Park

**Author notes:** Co-first author (Byung Hun Lee, Jae Youn Shim).

## Abstract

Memories are thought to be encoded in populations of neurons called memory trace or engram cells. However, little is known about the dynamics of these cells because of the difficulty in real-time monitoring of them over long periods of time *in vivo*. To overcome this limitation, we present a genetically-encoded RNA indicator (GERI) mouse for intravital chronic imaging of endogenous *Arc* mRNA—a popular marker for memory trace cells. We used our GERI to identify *Arc*-positive neurons in real time without the delay associated with reporter protein expression in conventional approaches. We found that the *Arc*-positive neuronal populations rapidly turned over within two days in the hippocampal CA1 region, whereas ∼4% of neurons in the retrosplenial cortex (RSC) consistently expressed *Arc* following contextual fear conditioning and repeated memory retrievals. Dual imaging of GERI and a calcium indicator in CA1 of mice navigating a virtual reality environment revealed that only the population of neurons expressing *Arc* during both encoding and retrieval exhibited relatively high calcium activity in a context-specific manner. This *in vivo* RNA imaging approach opens the possibility of unraveling the dynamics of the neuronal population underlying various learning and memory processes.

**One Sentence Summary:** Live-animal imaging of *Arc* mRNA reveals the dynamics and activity of memory trace cells during memory encoding and retrieval.

---

Activity-dependent gene expression is critical for long-term memory formation (*1*). Because immediate-early genes (IEGs) such as *Arc, c-Fos*, and *Egr-1* are rapidly and transiently transcribed within a few minutes following stimulation (*2*), their expression is widely used to identify neurons activated by diverse learning and memory tasks (*3*). Recent optogenetic and chemogenetic studies have demonstrated that IEG-expressing neurons are involved in formation of so-called memory traces or engrams (*4, 5*). Thus, IEG-positive neurons are often referred to as engram cells, which are defined by their activation for memory encoding and retrieval (*6*). However, the overlap between neuronal populations expressing IEGs during encoding and retrieval is relatively low, raising important questions as to whether technical limitations hinder precise identification of engram cells or whether engrams have an inherently dynamic nature (*7*).

Current methods for imaging IEG expression include RNA fluorescence *in situ* hybridization (FISH) (*2*) or immunostaining, and IEG promoter-driven expression of exogenous reporter proteins (*4, 8-18*). These methods have been used to take snapshots of the neuronal populations activated during memory encoding and retrieval. To date, however, it has not been possible to continuously monitor IEG transcription in real time in live animals because reporter proteins are typically expressed with a delay longer than an hour after transcription (*8, 9, 11, 13, 19*). For this reason, it is still largely unknown what type of neuronal activity triggers IEG transcription at the single-cell level. Moreover, even short-lived versions of fluorescent proteins decay over several hours to a few days (*8, 9, 15, 20*). Therefore, it has been difficult to identify distinct IEG-positive neuronal populations that are activated by different behaviors or events at a time interval less than a day. Various methods using tTA-TetO or tamoxifen-inducible Cre recombinase systems have been used to restrict IEG activity-tagging to a particular time window (*4, 10, 12, 14, 16, 18*), but the duration of the tagging window is typically hours to days. Hence, there have been concerns that these methods may overestimate the size of the neuronal population activated by a single experience (*7, 21*).

Herein, we report a genetically-encoded RNA indicator (GERI) imaging technique that enables the real-time monitoring of endogenous IEG transcription in the living brain. We generated a GERI mouse in which every single endogenous *Arc* mRNA was labeled with up to 48 green fluorescent proteins (GFPs) and visualized individual *Arc* transcription sites *in vivo* using two-photon excitation microscopy. By performing *in vivo* time-lapse imaging and simulations, we demonstrate that GERI reports the rapid and transient transcription of *Arc* mRNA with a high accuracy in real time, which has not been achieved by existing fluorescent protein reporters. Using this unique tool, we found that the *Arc*-positive neuronal populations have distinct dynamic properties in different brain regions. We also demonstrate that GERI can be used in conjunction with a genetically-encoded calcium indicator (GECI) to investigate the neuronal activity of *Arc*-positive neurons in awake mice navigating a virtual reality environment. This work establishes GERI as a versatile tool for probing the endogenous transcriptional activity during learning and memory processes *in vivo*.

## RESULTS

### Characterization of GERI mice for *in vivo* imaging of *Arc* mRNA

To label endogenous *Arc* mRNA with GFP, we exploited the highly specific binding between the PP7 bacteriophage capsid protein (PCP) and the PP7 binding site (PBS) RNA stem-loop (*22*). We generated a transgenic mouse that expresses a tandem PCP fused with tandem GFP (PCP-GFP) in neurons. This PCP-GFP mouse was crossed with an *Arc*-PBS mouse, in which 24 PBS repeats were knocked into the 3’ untranslated region (UTR) of the *Arc* gene (*23*). In the resulting PBS homozygous (*Arc*^P/P^) PCP×PBS hybrid mouse (Figure 1A), every endogenous *Arc* mRNA was labeled with up to 48 GFPs. To confirm that PP7-GFP labeling did not disrupt *Arc* gene expression, we performed Western blotting and immunofluorescence analyses to evaluate the Arc protein expression levels in the brains of wild-type (WT), *Arc*-PBS, and PCP×PBS mice (Figure S1), and the results indicated that the Arc protein levels were similar in all mice.

**Figure 1.**
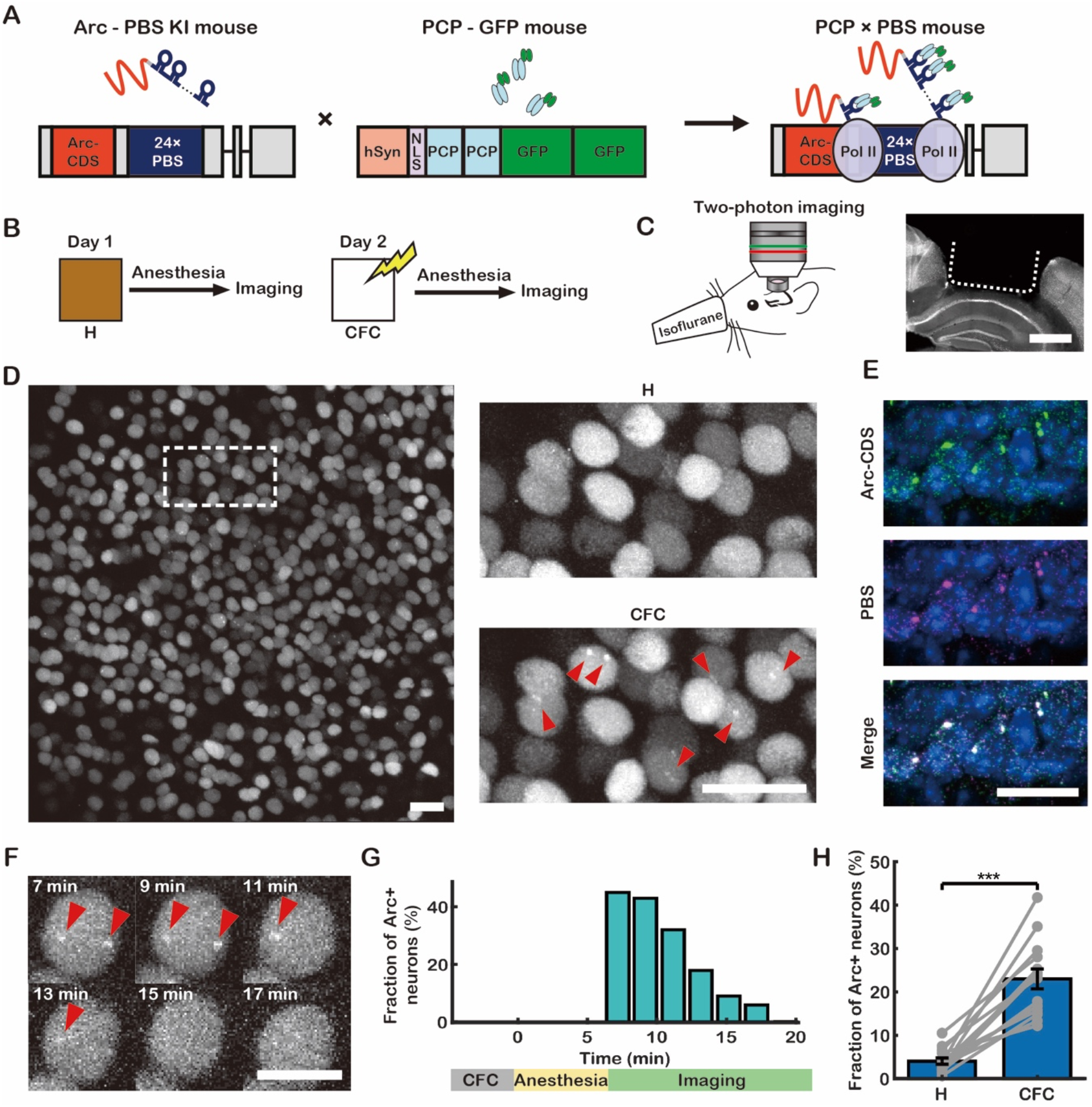
Development of genetically encoded RNA indicator (GERI) mice for imaging *Arc* mRNA. **(A)** Schematic for labeling *Arc* mRNA *in vivo*. NLS, nuclear localization sequence; Pol II, RNA polymerase II. (Gray boxes, UTR; red box, *Arc* coding sequence (*Arc*-CDS); blue box, 24× PBS cassette; black lines, introns). The *Arc*-PBS knock-in (KI) mouse was crossed with the PCP-GFP mouse to generate PCP×PBS hybrids. **(B)** On day 1, mice were removed from their home cage (H) and immediately anesthetized for *in vivo* imaging. On day 2, the mice were subjected to contextual fear conditioning (CFC) followed by *in vivo* imaging. **(C)** Left, experimental setup for *in vivo* two-photon imaging through a hippocampal window. Right, coronal view of the brain of a PCP×PBS mouse after hippocampal window surgery. **(D)** Representative *in vivo* image of CA1 neurons in a PCP×PBS mouse after autofluorescence subtraction (for detailed image processing procedures, see Figure S2). The same region (dotted box) is enlarged for comparison of images taken on day 1 (H) and day 2 (CFC). *Arc* transcription sites are marked with red arrowheads. **(E)** *Arc* mRNA detected by two-color smFISH targeting *Arc*-CDS (green) and PBS (magenta). **(F)** Time-lapse images of an *Arc*+ neuron after CFC. **(G)** Fraction of *Arc*+ neurons over time after CFC. **(H)** Fraction of *Arc*+ neurons in CA1 after H and CFC conditions (*n* = 12 mice; *** *P* < 10^−4^ by pairwise *t* test). Scale bars, (C) 1 mm, (D and E) 50 μm, and (F) 10 μm. Error bars represent the standard error of the mean (SEM).

To demonstrate that activity-dependent *Arc* expression can be observed by *in vivo* imaging of PCP×PBS mice, we first investigated *Arc* mRNA induction upon contextual fear conditioning (CFC) (Figure 1B). We used two-photon excitation microscopy through a hippocampal window (*24*) to visualize *Arc* mRNA in the dorsal CA1 region of the PCP×PBS mice (Figure 1C). To minimize motion artifacts and any effects of subsequent handling, we performed *in vivo* imaging of mice under anesthesia immediately after CFC; previous reports have demonstrated that anesthesia with isoflurane does not cause retrograde amnesia (*25, 26*). We developed image processing algorithms for motion correction, autofluorescence subtraction, region of interest (ROI) registration, and automatic segmentation of neurons by their nuclear GFP background originating from the nuclear localization sequence (NLS) in the PCP-GFP (Figure S2). Using the software, we were able to identify the same neurons across multiple days and weeks. In a subset of neurons, one or two bright spots were clearly visible in the nucleus (red arrowheads, Figure 1D). To confirm that these were *Arc* transcription sites, we performed two-color single-molecule FISH (smFISH) using probes targeting the coding sequence (CDS) and the PBS sequence of *Arc* mRNA in brain tissues collected after CFC (Figure 1E). *Arc* mRNA particles in smFISH images were detected and classified by their size and intensity into two major groups: single mRNAs and transcription sites. Approximately 93% of the transcription sites detected by the PBS probe were colocalized with those detected by the CDS probe, and vice versa (Figure S3). The average copy number of nascent *Arc* mRNA per transcription site was estimated to be 15 ± 4 by dividing the intensity of each transcription site by the average intensity of single mRNAs detected by the PBS probe. Thus, an average of ∼720 GFPs were recruited to each *Arc* locus, providing sufficient signal for three-dimensional visualization of individual transcription sites located 100−400 μm deep inside the living brain (Movie S1).

To investigate the temporal dynamics of *Arc* transcription, we performed time-lapse imaging and found that the fraction of neurons with *Arc* transcription sites (*Arc*+ neurons) reached its maximum within 7 min after CFC and then monotonically decreased to basal levels by 20 min after CFC (Figures 1F and 1G). Based on these data, the time window selected for *in vivo* imaging was 4–7 min after each behavioral test, substantially faster than the 1−3 hours after stimulation for existing fluorescent protein reporters (*8, 9, 13, 19*). To evaluate the performance of GERI compared with a short half-life GFP (shGFP) reporter, we performed Monte-Carlo simulations and assessed the accuracy of classifying IEG-positive and IEG-negative cells. The simulation results suggested that the classification accuracy of GERI became superior to that of the shGFP reporter as the basal IEG activation rate was increased (Figure S4).

Having established GERI imaging and analysis techniques, we measured the size of the *Arc*+ neuronal populations in CA1 of the PCP×PBS mice. We found *Arc* transcription sites in 4.1 ± 0.8% of the CA1 neurons in mice taken from their home cages (H). The percentage of *Arc*+ neurons significantly increased to 23 ± 3% in the same ROIs after CFC on the next day (Figures 1D and 1H). Three-color smFISH showed that *Arc* and two other IEGs, *c-Fos* and *Egr-1*, were regulated similarly and coexpressed in 19 ± 3% of the neurons in the dorsal CA1 region after CFC (Figure S5). In addition, we found that *Arc* transcription sites were readily detected by GERI without any staining in fixed sections of various brain regions (Figure S6), which provides another application of GERI for RNA *in situ* detection.

### Dynamics of the *Arc*+ neuronal population in recent memory retrieval

Using our *in vivo* GERI imaging technique, we next investigated how the populations of *Arc*+ neurons change upon memory encoding and retrieval. We performed fear conditioning in context A (ctx A) and repeatedly exposed the mice to ctx A to retrieve fear memory on three consecutive days (Figure 2A). The animals showed freezing behavior during all retrieval tests (R1–3) in ctx A but not in a new context (ctx B) (Figure 2B), indicating that conditioned context-specific fear memory remained intact. To compare the dynamics of the *Arc*+ neuronal population in different brain regions, we performed cranial window imaging of the dorsal hippocampal CA1 region and the retrosplenial cortex (RSC), which have essential roles in spatial cognition and memory with strong reciprocal connectivity (*27-29*). A similar fraction (14–20%) of neurons in CA1 transcribed *Arc* after CFC and R1–3 (Figure 2C). The fraction of *Arc*+ neurons in the RSC was higher than that in CA1 and increased from 24 ± 4% (CFC) to 35 ± 7% (R3) (Figure 2D).

**Figure 2.**
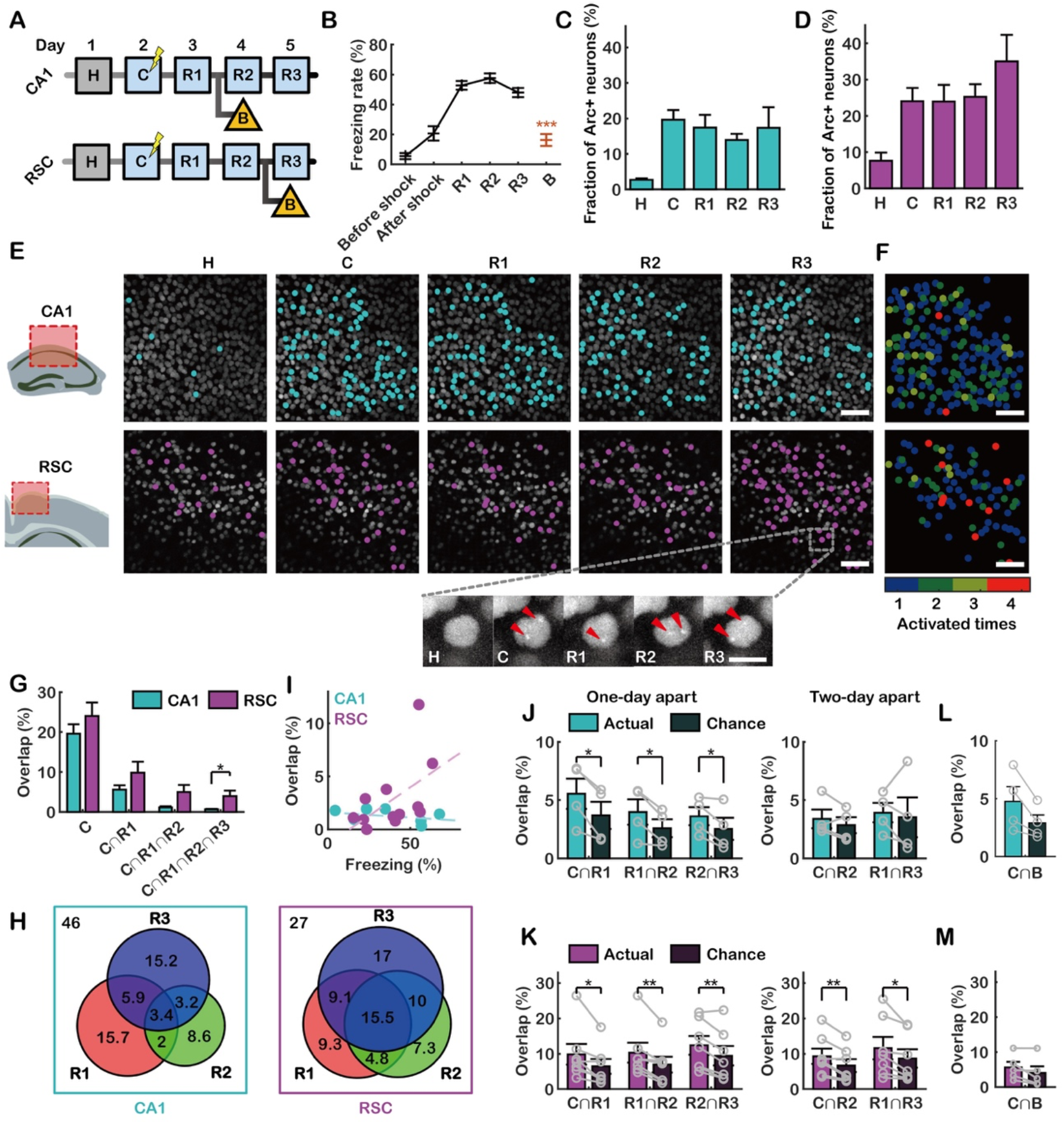
Dynamics of *Arc*+ neuronal populations following CFC and repeated recent memory retrieval. **(A)** Experimental scheme for CFC (C) and repeated retrieval tests (R1-3) over three consecutive days. Groups of mice were exposed to context B on day 4 (CA1) or day 5 (RSC). **(B)** Freezing rates of PCP×PBS mice during CFC and three retrieval tests in ctx A (*n* = 11 mice) and ctx B (*n* = 9 mice) (Tukey’s multiple comparison test after one-way ANOVA, *** *P* < 10^−6^). **(C-D)** Fraction of *Arc*+ neurons in CA1 (C) (*n* = 4 mice) and the RSC (D) (*n* = 7 mice) after each behavioral session. **(E)** Representative *in vivo* images of CA1 (upper panels) and the RSC (lower panels), showing the same fields of view after H, C, and R1–3. Cyan and magenta dots denote *Arc*+ neurons in CA1 and the RSC, respectively. Bottom, example images of a neuron with red arrows indicating *Arc* transcription sites. (**F**) *Arc*+ neurons in CA1 (top) and the RSC (bottom), colored by the number of times the neurons expressed *Arc*. **(G)** Overlap percentage of consecutively reactivated *Arc*+ neurons in CA1 and the RSC (* *P* < 0.05 by rank-sum test). **(H)** Venn diagrams of neurons that expressed *Arc* during CFC and re-expressed *Arc* during each retrieval test (left: CA1, right: RSC). Numbers indicate percentages of neurons. **(I)** The freezing rate was correlated with the overlap rate of *Arc*+ neurons in the RSC (magenta) but not in CA1 (cyan) (CA1: *R* = -0.40, *P* = 0.84, *n* = 8 mice; RSC: *R* = 0.52, *P* = 0.04, *n* = 12 mice; by Pearson’s correlation). **(J-K)** Overlap percentage of *Arc*+ neurons at one- (left) or two-day intervals (right) (* *P* < 0.05, ** *P* < 0.01 by pairwise *t* test) in CA1 (J) and the RSC (K). **(L-M)** Overlap of *Arc*+ populations between the CFC and ctx B conditions compared with chance in CA1 (L; *P* = 0.055) and the RSC (M; *P* = 0.12). Scale bars, (E and F) 50 μm and (E, inset) 10 μm. Error bars represent the SEM.

Taking advantage of our live animal imaging, we performed longitudinal analysis of *Arc* transcription in individual neurons. We found that a distinct population of neurons expressed *Arc* mRNA every time contextual memory was retrieved (Figure 2E). We then identified the overlapping population of *Arc*+ neurons across the CFC and R1–3 (Figure S7). A small population of neurons persistently showed *Arc* transcription throughout both the conditioning and retrieval sessions (Figure 2E, bottom panel). This persistently overlapping population (red dots, Figure 2F) represented 4.0 ± 0.2% of the neurons in the RSC, which was significantly higher than that in CA1 (0.6 ± 0.1%) (Figure 2G). The population of neurons that expressed *Arc* after CFC and each retrieval session also overlapped more in the RSC than in CA1 (Figure 2H). Moreover, we found a positive correlation between the freezing rate and the fraction of the persistently overlapping *Arc*+ population in the RSC, whereas there was no such correlation in CA1 (Figure 2I). Since the freezing rate reflects the strength of fear memory, the persistently overlapping *Arc*+ neurons in the RSC could be a stable component of contextual fear memory (*10*). However, there were more dynamic changes occurring in the *Arc*+ neuronal population in CA1 during memory encoding and retrieval.

To assess the dynamics of *Arc* re-expression, we compared the overlap of *Arc*+ neurons upon re-exposure to ctx A at one- or two-day intervals to that expected by chance. In CA1, the overlap was greater than expected by chance in the one-day-apart comparison, but equivalent to random in the two-day-apart comparison (Figure 2J). In the RSC, the overlap in both comparisons was significantly greater than that expected by chance (Figure 2K). We confirmed that the degree of overlap between the *Arc*+ neurons activated during CFC and those activated upon exposure to ctx B was almost random in both CA1 and the RSC (Figures 2L and 2M). These data indicate that there was a near complete turnover of *Arc*+ neurons within two days in CA1, consistent with previous reports (*15, 30-32*). In contrast, the *Arc*+ neurons in the RSC exhibited either slower dynamics or greater stability.

### Stability of the overlapping *Arc*+ population in remote memory retrieval

To examine the stability of *Arc*+ neuronal ensembles over longer time scales, we performed repeated retrieval experiments on days 9, 16, 23, and 30 (Figure 3A). All fear-conditioned mice showed freezing behaviors during the remote memory retrieval tests (Figure 3B). The fraction of *Arc*+ neurons in CA1 decreased slightly over a month (Figure 3C), whereas the fraction in the RSC was similar (28–34%) for all sessions (Figure 3D). The overlapping population of *Arc*+ neurons sharply decreased in CA1 but remained stable in the RSC for at least one month (Figure 3E). Persistent *Arc* re-expression up to the fourth retrieval session was observed in 4.0 ± 1.3% of the RSC neurons but in almost none (0.2 ± 0.2%) of the CA1 neurons (Figure 3F). The consecutive overlap of *Arc*+ neurons was significantly higher in the RSC than in CA1 from R2 (Figure 3G). The overlap between the *Arc*+ populations elicited by memory retrieval 1–4 weeks apart was also mostly higher than that expected by chance in the RSC but not in CA1 (Figure S8). To visualize how *Arc*+ neurons turn over during remote memory retrieval, we used tree graphs to trace *Arc* re-expression probabilities (Figure 3H). RSC neurons that consistently expressed *Arc* were more likely to be reactivated, in accordance with previous reports on neocortical IEG expression (*9, 19, 27, 33*). However, in CA1, an almost random population of neurons expressed *Arc* whenever the mouse was re-exposed to the same context.

**Figure 3.**
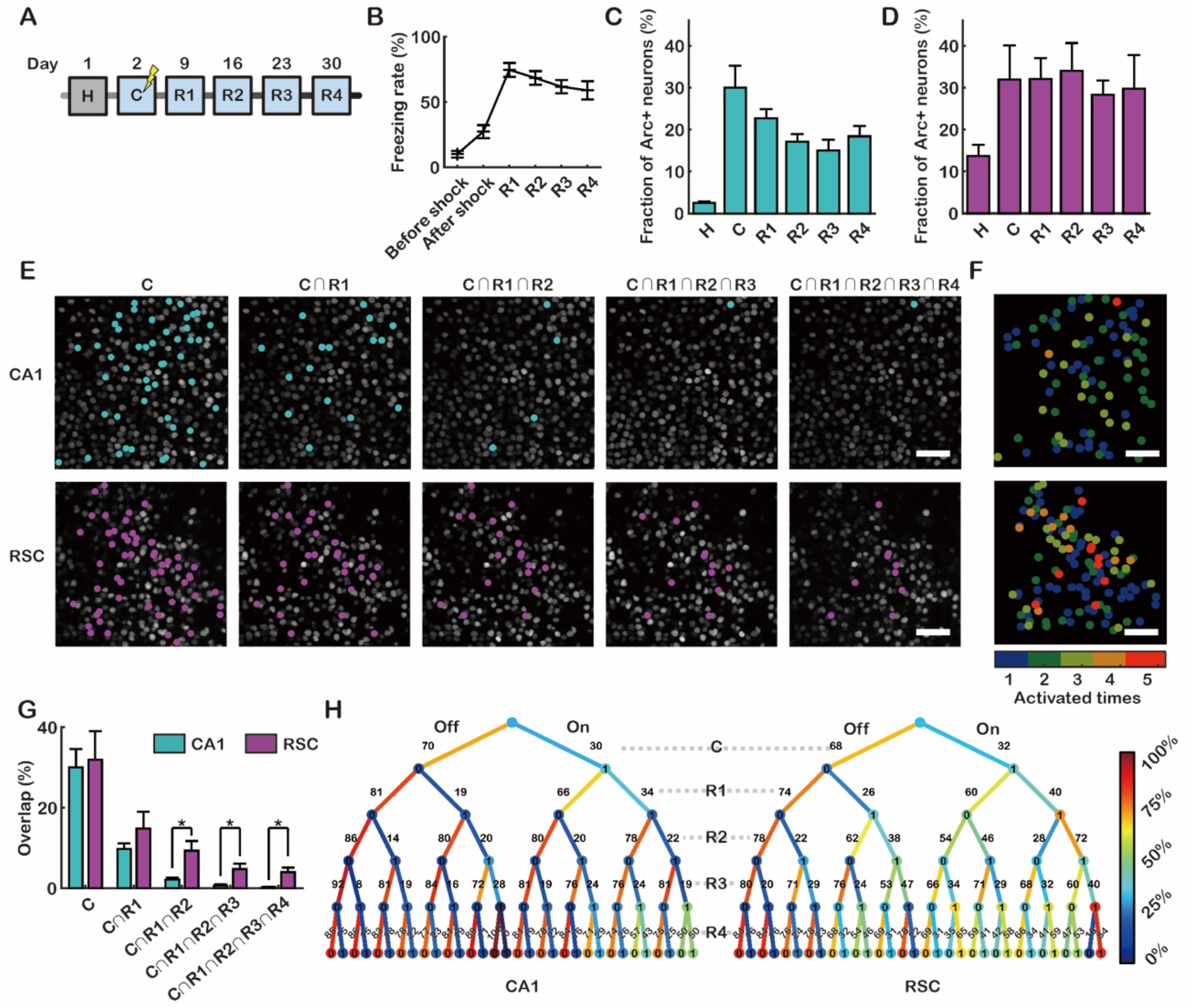
Long-term dynamics of *Arc*+ neuronal ensembles in CA1 and the RSC. **(A)** Experimental scheme for CFC (C) and remote memory retrieval tests (R1−4). **(B)** Freezing rates of PCP×PBS mice during CFC and four retrieval tests (*n* = 8 mice). **(C-D)** Fraction of *Arc*+ neurons in CA1 (C) (*n* = 4) and the RSC (D) (*n* = 4) after each session. (**E**) Representative images of CA1 (top) and the RSC (bottom) showing overlapping populations of *Arc*+ neurons (cyan, CA1; magenta, RSC). **(F)** *Arc*+ neurons in CA1 (top) and the RSC (bottom) labeled to reflect the number of *Arc* expression events. **(G)** The percentage of neurons that consecutively re-expressed *Arc* in CA1 (cyan) and the RSC (magenta) (* *P* < 0.05 by Student’s t test). (**H**) Tree graphs colored by the re-expression probability of neurons in CA1 (left) and the RSC (right). The first layer represents the fraction of neurons that did (rightwards) or did not express (leftwards) *Arc* following CFC. The other layers are further split according to the presence or absence of *Arc* transcription sites following memory retrieval. Scale bars, (E and F) 50 μm. Error bars represent the SEM.

### Calcium activity of *Arc*+ neurons during VR navigation

Given our data on the high turnover rate of *Arc*+ neurons in CA1, we next sought to investigate the neuronal activity of these cells during memory encoding and retrieval. To monitor the calcium activity of *Arc*+ neurons, we performed dual-color imaging of CA1 in head-fixed awake mice running on a trackball to navigate a linear track in virtual reality (VR) (Figure 4A). We injected adeno-associated virus (AAV) expressing jRGECO1a (*34*) into the dorsal CA1 region of the PCP×PBS mice (Figure S9). Six mice were exposed to a novel context (ctx A) on day 1 and again on day 2, followed by a distinct context (ctx B) on day 3 (Figure 4B). Each day, we imaged *Arc* mRNA in the awake resting state, calcium activity during VR navigation, and *Arc* mRNA under anesthesia (Figures 4C and S10, and Movie S2). The fraction of neurons with *Arc* transcription sites significantly increased after VR navigation each day (Figure 4D). The calcium activity of the *Arc*+ neurons was similar regardless of the prior transcription of *Arc* before VR navigation (Figure S11). Cells that transcribed *Arc* after VR navigation on days 1, 2, and 3 were referred to as A1-*Arc*+, A2-*Arc*+, and B-*Arc*+ neurons, respectively; the overlap between the three *Arc*+ populations was 15–17% (Figure 4E)—significantly greater than expected by chance (Figure S12). We next compared the calcium activity of the *Arc*+ and *Arc*− neurons pooled from all three sessions. The calcium traces were deconvolved to infer spike trains (*35*), from which we calculated the ‘inferred burst’ and ‘inferred theta-burst’ (6– 10 Hz) rates (Figure S11). By applying the same analysis to cultured neurons under electrical stimulation, we confirmed that burst activity at a frequency of 6–10 Hz could be detected by using this deconvolution algorithm (Figure S13). Although almost all neurons (99.9%) showed varying degrees of calcium activity (Figure 4F), we found that *Arc*+ neurons were significantly more active than *Arc*− neurons on average, in terms of calcium event, inferred burst, and inferred theta-burst rates (Figures 4G and S11).

**Figure 4.**
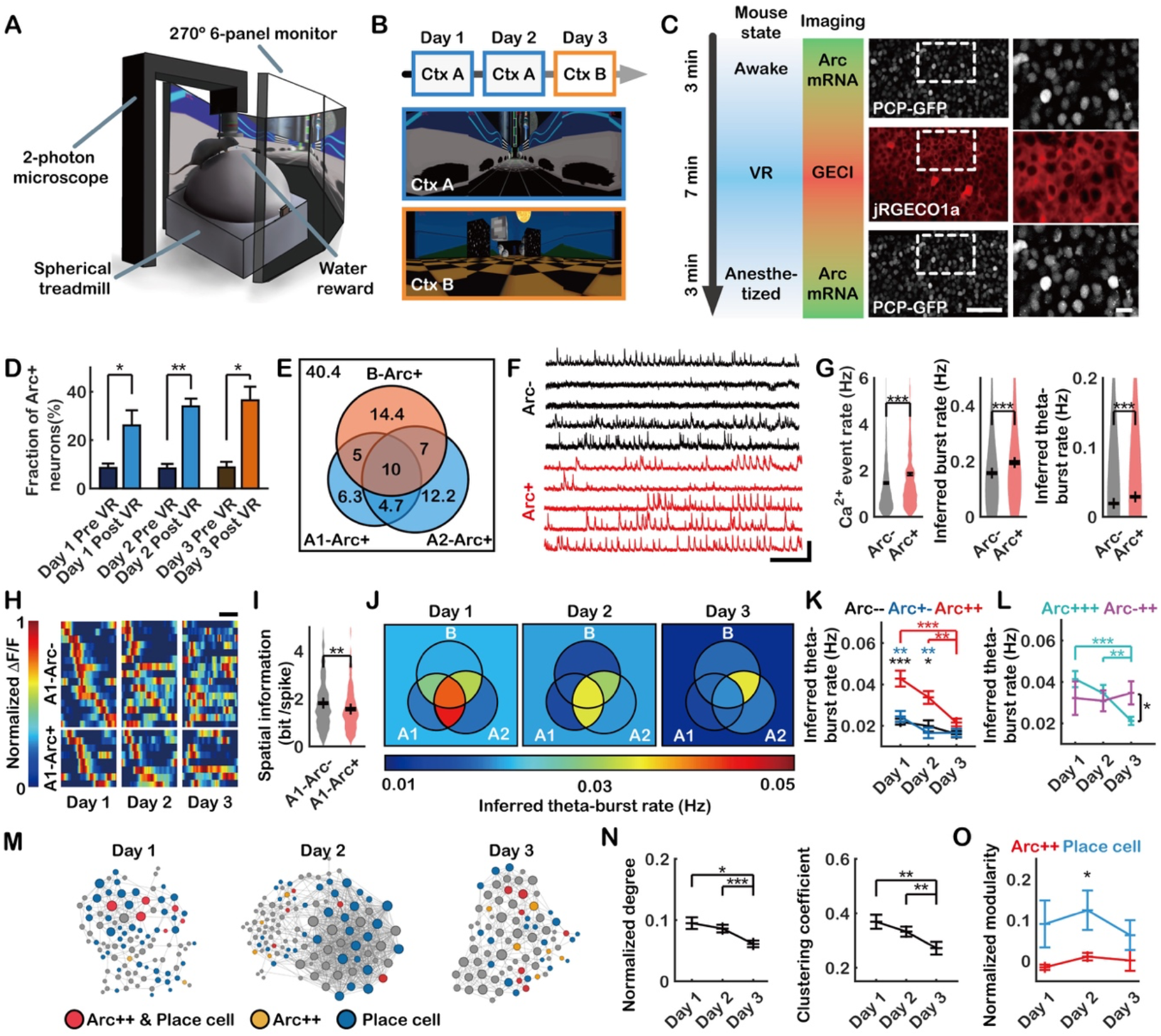
*Arc* transcription and calcium imaging in CA1 of mice exploring virtual reality (VR). **(A)** Experimental setup. **(B)** Experimental schedule (top) and virtual contexts A (middle) and B (bottom). **(C)** *Arc* transcription imaging was first conducted while the mice were awake, followed by calcium imaging while exploring VR. After 7 min of exploration, *Arc* transcription imaging was performed under anesthesia. Representative images and magnified insets shown on the right. **(D)** Fractions of neurons with *Arc* transcription sites before and after VR on each day (* *P* < 0.05, ** *P* < 0.01 by pairwise *t* test, *n* = 6 mice). **(E)** A Venn diagram of *Arc*+ neurons on days 1, 2, and 3, showing the percentage of each subpopulation. **(F)** Representative calcium traces of *Arc*− and *Arc*+ neurons. **(G)** Left panel, Ca^2+^ event rates of *Arc*− and *Arc*+ neurons (*** *P* < 10^−8^ by rank-sum test). Middle, inferred burst rates of *Arc*− and *Arc*+ neurons (*** *P* < 10^−10^ by rank-sum test). Right panel, inferred theta-burst rates of *Arc*− and *Arc*+ neurons (*** *P* < 10^−19^ by rank-sum test; *Arc*−: *n* = 1085 neurons, *Arc*+: *n* = 651 neurons). **(H)** Example of place fields of A1-*Arc*− (top) or A1-*Arc*+ (bottom) place cells on days 1, 2 and 3. Neurons are ordered by the locations of the place field that yielded the maximum activity on day 1. **(I)** Spatial information of A1-*Arc*− and A1-*Arc*+ neurons on day 1 (** *P* < 0.01 by rank-sum test; A1-*Arc*−: *n* = 430 neurons, A1-*Arc*+: *n* = 209 neurons). **(J)** Venn diagrams of average *Arc*+ neuron inferred theta-burst rates on days 1, 2 and 3. **(K)** Inferred theta-burst rates of *Arc*−−, *Arc*+− and *Arc*++ neurons on each day (* *P* < 0.05, ** *P* < 0.01, *** *P* < 10^−4^ by Student’s *t* test; *Arc*−−: *n* = 261−302 neurons, *Arc*+−: *n* = 69−92 neurons, *Arc*++: *n* = 103−116 neurons). **(L)** Inferred theta-burst rate of *Arc*−++ and *Arc*+++ neurons on each day (* *P* < 0.05, ** *P* < 0.01, *** *P* < 10^−4^ by Student’s *t* test; *Arc*−++: *n* = 36−47 neurons, *Arc*+++: *n* = 65−81 neurons). **(M)** Example network graphs from a mouse on days 1, 2 and 3. Nodes are classified by color, and node size is proportional to the degree of the neurons. **(N)** Normalized degree (left) and clustering coefficient (right) of *Arc*++ neurons on each day (* *P* < 0.05, ** *P* < 0.01, *** *P* <10^−4^ by rank-sum test; *n* = 103−116 neurons). **(O)** Normalized modularity when dividing neurons into place cells and nonplace cells (blue) and dividing neurons into *Arc*++ and non-*Arc*++ (red) (* *P* < 0.05 by Student’s *t* test; *n* = 6 mice). Scale bars, (C, middle) 50 μm, (C, right) 10 μm, (F, horizontal) 1 min, (F, vertical) 150% ΔF/F and (H) 1 m. Error bars represent the SEM.

Next, we identified place cells on each day using a similar method previously reported (*24*) and calculated the spatial correlation between the corresponding place fields (Figures 4H and S14A-C). The correlation between place fields in the same context (A-A) was significantly higher than that in different contexts (A-B), indicating that the mice perceived the two virtual environments as different contexts (Figure S14D). Among the place cells, A1-*Arc*+ neurons showed weaker spatial correlations between days 1 and 2 than A1-*Ar*c− neurons (Figure S14E). A1-*Arc*+ neurons also had lower spatial information than A1-*Arc*− neurons on day 1 (Figure 4I), consistent with a previous report on *c-Fos*-positive neurons (*36*). These results support the notion that IEG-positive CA1 neurons do not necessarily represent spatial information about the context, but rather serve as an index to episodic memory (*37*).

To further dissect the activity of *Arc*+ neurons, we grouped the CA1 neurons into eight subpopulations, as shown in Figure 4E, and compared their calcium activity on each day (Figures 4J and S15A). Neurons that transcribed *Arc* on both days 1 and 2 (*Arc*++), which includes both *Arc*+++ and *Arc*++− neurons, had higher inferred theta-burst activity in ctx A than in ctx B (Figure 4K). Neurons that expressed *Arc* on day 1 but not on day 2 (*Arc*+−) showed a similar inferred theta-burst rate as neurons that did not express *Arc* on either day (*Arc*−−) throughout days 1 to 3 (Figure 4K). Both *Arc*+− and *Arc*−− neurons exhibited significantly lower inferred theta-burst activity than *Arc*++ neurons on days 1 and 2 (Figure 4K). These data suggest that only *Arc*++ neurons are involved in contextual memory for ctx A through relatively high theta-burst activity during both encoding and retrieval. Interestingly, neurons that expressed *Arc* on all three days (*Arc*+++) also showed significantly higher inferred theta-burst rates in ctx A than in ctx B (Figures 4J and 4L), indicating that these neurons are involved in the memory for ctx A but not for ctx B. On the other hand, neurons that expressed *Arc* only on days 2 and 3 (*Arc*−++) showed a high inferred theta-burst rate on day 3 (Figures 4J and 4L). These results suggest that only a subpopulation of IEG-positive neurons (such as *Arc*++ neurons) support memory for a specific context.

We then examined whether these overlapping *Arc*++ neurons were simultaneously reactivated during memory retrieval as in optogenetic experiments (*37*). The correlation coefficients among the *Arc*++ neurons were similar to those among the non-*Arc*++ neurons (Figures S15B and S15C). After calculating the correlation coefficient matrices, we generated network graphs of the correlated neuronal activity on each day (Figure 4M). The network properties were quantified using the number of correlated pairs (degree), the fraction of pairs among neighbors (clustering coefficient) (*38*), and the strength of division of the network into modules (modularity) (*39*). *Arc*++ neurons showed a higher normalized degree and clustering coefficient in ctx A than ctx B (Figure 4N), indicating that their activity was highly correlated with that of other CA1 neurons during encoding and retrieval of contextual memory. The modularity of *Arc*++ neurons was close to zero (Figure 4O), suggesting that they were integrated with rather than segregated from other neurons in CA1. Meanwhile, the place cells identified each day showed higher modularity than the *Arc*++ neurons (Figure 4O) and formed a dense cluster on day 2 (Figure 4M), consistent with a previous report (*40*). These results suggest that *Arc*++ neurons do not necessarily have synchronized inputs but rather exhibit correlated activity with other CA1 neurons in a context-specific manner.

## Discussion

Our GERI imaging technique enables the real-time visualization of endogenous IEG transcription at individual gene loci in live animals. Unlike conventional reporter protein expression approaches, GERI can directly report the rapid and transient transcription of IEGs with high accuracy. Using this unique tool, we found that the *Arc*+ neuronal populations in the hippocampus and the cortex have distinct dynamic properties. In CA1, a similar fraction (∼20%) of neurons expressed *Arc* during both recent and remote memory retrieval, but the individual neurons involved rapidly changed within two days (Figure 2J). However, in the RSC, a small yet significant fraction (∼4%) of neurons persistently expressed *Arc* during encoding and in every retrieval session for at least one month (Figure 3G). The proportion of this persistent *Arc+* population in the RSC was correlated with the freezing rate (Figure 2I), indicating a role for these stable *Arc+* neurons in contextual fear memory. These findings are particularly interesting in the context of memory indexing theory, in which hippocampal ensembles are proposed to serve as indices, whereas cortical ensembles serve as content (*41*). From this perspective, our results suggest that each hippocampal *Arc*+ population represents a new or updated index for each retrieval event, while the stable RSC *Arc*+ population contains information about context (*37*). Moreover, the drastic decrease in the overlapping *Arc*+ population in CA1 implies that the role of the hippocampus in systems memory consolidation is limited to short time scales of a few days (*6, 42, 43*).

Although IEG expression has long been used as a marker for recent activity (*2*), the exact relationship between neuronal activity and IEG transcription at the single-cell level has not been established. By combining GERI with GECI imaging in awake mice navigating a VR environment, we found that the *Arc*+ population had higher average calcium activity than the *Arc*− population, but individual neurons in both groups showed widely varying degrees of calcium activity (Figures 4F and 4G). Only the overlapping *Arc*++ subpopulation in CA1 consistently exhibited higher inferred theta-burst activity than other neurons during both memory encoding and retrieval (Figures 4J and 4K), suggesting that the *Arc*++ population, rather than the whole *Arc*+ population, may be involved in contextual memory.

Because *Arc* is a key regulator of synaptic plasticity (*44*), *Arc* expression during memory retrieval may have a role in modifying or updating the representation of memory. From this perspective, an alternative interpretation is possible, in which the neurons that express *Arc* during encoding but not during retrieval (*Arc*+−) could be the neuronal population that maintains the original memory. However, we found that *Arc*+− neurons showed similar levels of calcium activity as neurons that never expressed *Arc* (*Arc*−−) during exposure to either ctx A or ctx B (Figures 4J and 4K). Therefore, our data support the interpretation that the overlapping *Ar*c++ subpopulation is more likely to be involved in the so-called memory trace or engram. Further technical developments for the selective manipulation of subpopulations such as *Ar*c++ would be necessary to more precisely identify memory trace or engram cells.

The live-animal RNA imaging approach demonstrated here provides a powerful platform for future experiments to elucidate the dynamics of memory formation in conjunction with various calcium or voltage indicators, biosensors (*45*), and optogenetic tools (*46*). Moreover, different GERI tools could be designed using newly developed background-free RNA labeling systems such as MS2-PP7 (*47, 48*) to achieve a higher signal-to-noise ratio for RNA imaging in freely behaving mice. Ultimately, we expect GERI to enable visualization of even single RNA molecules in subcellular compartments such as dendritic spines for studies of RNA localization (*23, 49, 50*) in the live brain.

## Acknowledgements

We thank D. A. Dombeck for transferring the hippocampal window surgery technique; B. K. Kaang and H. Kim for their advice on behavioral experiments and helpful discussions; I. Lee and E.-H. Park for sharing their expertise in VR experiments; and S. Kim and M. Kim for mouse colony management and assistance with image analysis. This research was supported by the Samsung Science and Technology Foundation under Project Number SSTF-BA1602-11. The VR equipment was acquired with funding from the Howard Hughes Medical Institute– Wellcome International Research Scholar Award from the Wellcome Trust (208468/Z/17/Z).

## Author contributions

J.Y.S. and H.Y.P. developed the PCP-GFP mouse model. B.H.L. and J.Y.S. performed CFC and *in vivo* imaging of *Arc* transcription. B.H.L. and D.W.K. performed VR experiments with *in vivo* imaging of calcium activity and *Arc* transcription. B.H.L. developed the image-processing algorithms. B.H.L., J.Y.S., H.C.M., and D.W.K. performed the image analysis. H.C.M. and J.Y.S. performed the Western blot analysis. J.K., J.S.Y. and J.K. performed smFISH, immunofluorescence, and GFP imaging in fixed brain slices. B.H.L. and H.Y.P. wrote the paper. H.Y.P. conceived and supervised the project.

## Data and materials availability

All materials generated in this study are available from the lead contact with a completed material transfer agreement.

