## Supplementary information for "Real-time visualization of mRNA synthesis during memory formation in live animals"

**This PDF file includes:**

Materials and Methods

Figs. S1 to S15

Captions for movies S1 and S2

References

**Other supplementary materials for this manuscript includes the following:**

Movies S1 and S2

### Materials and Methods

Generation of PCP-GFP transgenic mouse line

Animal care practices and all experimental protocols were approved by the Institutional Animal Care and Use Committee at Seoul National University. We generated a transgenic mouse line that expresses a synonymous tandem PP7 capsid protein fused with tandem GFP (stdPCP-stdGFP) (*1*) under the control of the human synapsin-1 promoter (*hSyn*). The *hSyn-stdPCP-stdGFP* transgene was flanked by two *Rosa26* arms, and the resulting DNA fragments were microinjected into zygotes from C57BL/6N wild type (WT) mice. The transgene integration site was found between positions 26,509,464 and 26,536,787 on chromosome 13 by whole genome sequencing.

Genotyping

Genotyping of PCP-GFP transgenic mice was performed by PCR analysis using the following primer sets. For the PCP-GFP allele, we used forward primer hSyn_F (5′-CGACTCAGCGCTGCCTCAGTCT-3′) and reverse primer PCP_R (5′-CGTGTATCTAACCTTAGGTAGACC-3′), yielding a 383-bp product. For the WT allele, we used forward primer Ch13_F (5′-GTCTAGAGTGCTGCTTGTCTCC-3′) and reverse primer Ch13_R (5′-CTGTGCTTCAAAACCCCATGACC-3′), yielding a 733-bp product. PCR was conducted at 95 °C for 30 sec, 60 °C for 30 sec, and 72 °C for 60 sec for 35 cycles.

PCP-GFP and Arc-PBS knock-in mice were cross-bred (*2*) to obtain a double homozygous PCP×PBS hybrid mouse line. For genotyping of the Arc-PBS knock-in, we performed PCR analysis using the following primer sets. For the 5′ end, we used two forward primers, Arc PBS gt 5F (5′-TGTCCAGCCAGACATCTACT-3′) and ArcPBS gt 5R (5′-

TAGCATCTGCCCTAGGATGT-3′), and one reverse primer, PBS scr R1 (5′-GTTTCTAGAGTCGACCTGCA -3′), yielding a 320-bp product for the WT *Arc* allele and a 228-bp product for the PBS knock-in allele. For the 3′ end, we used forward primer Arc PBS gt 3F (5′-GACCCATACTCATTTGGCTG-3′) and reverse primer ArcPBS gt 3R (5′-GCCGAGGATTCTAGACTTAG-3′), yielding a 332-bp product for the WT *Arc* allele and a 413-bp product for the PBS knock-in allele. PCR was conducted at 94 °C 30 sec, 55 °C 30 sec, 72 °C 30 sec for 35 cycles.

Western blotting

Six-week-old female mice were housed in the home cage for 7 days and habituated by daily handling. On the eighth day, CFC was performed in Context A. After CFC, the mice were housed in the home cage for 1 hour and sacrificed by cervical dislocation after isoflurane anesthesia. Brain tissue extracts were prepared using T-PER tissue protein extraction reagent (Thermo Fisher Scientific, 78510) containing 1× protease inhibitor (Roche, 04693159001). 30 μg aliquots of protein were separated on 4-12% Bis-Tris polyacrylamide precast gels (Invitrogen, NW04120BOX) in MES-SDS running buffer (Invitrogen, B0002) and transferred to nitrocellulose membranes by using the Mini Blot Module (Thermo Fisher Scientific) following the manufacturer's instructions. The following antibodies were used: Anti-Arc (1:200, Santa Cruz Biotechnology, sc-17839) and anti-GAPDH (1:100,000, Sigma, G9545) as primary antibodies, and anti-rabbit IgG conjugated to HRP (1:5,000, SA002, GenDEPOT), and anti-mouse IgG conjugated to HRP (1:5,000, SA001, GenDEPOT) as secondary antibodies. Western blots were scanned by LAS 4000 (GE Healthcare Life Sciences). The images were analyzed by Image Studio Lite Ver 5.2 (LI-COR Biosciences).

Immunohistochemistry

12–13-week-old male mice were housed in the home cage and habituated by handling for 3 days. Then, the mice were subject to contextual fear conditioning and returned to the home cage for 1.2–1.4 hr. After isoflurane anesthesia, the animals were perfused transcardially with 15 ml of cold phosphate-buffered saline (PBS) containing 10 U/ml heparin (Sigma, H3393) for 3–4 min and 30 ml of cold 4% PFA in 0.1 M PBS for 7-8 min. Brains were post-fixed in 4% PFA in 0.1 M PBS at 4 °C for more than 4 hr and embedded in 3% agarose gel in 0.1 M PBS. With a vibratome (Leica, VT1000s), brains were sectioned coronally to 50 µm, permeabilized with 1% Triton X (Thermo Fisher Scientific, 28314) in Tris-buffered saline (TBS) for 30 min at room temperature (RT), and incubated in 0.4% Triton X and 5% normal goat serum (NGS) (Vector Labs, S-1000) in TBS for 30 min at RT. Sections were treated with primary antibodies in 0.4% Triton X and 5% NGS in TBS at 4 °C overnight and then washed three times with 0.1% Triton X in TBS for 20 min at RT. After secondary antibody treatment for 3 hr at RT, the sections were counterstained with 1 µg/ml DAPI (Thermo Fisher Scientific, 62248) in 0.1% Triton X in TBS. Then, the sections were washed with 0.1% Triton X in TBS for 20 min at RT and then twice with TBS for 20 min, and cover-slipped with VectaShield mounting media (Vector Labs, H-1700). Anti-Arc (1:1000, Synaptic Systems, 156 003) was used as primary antibody, and goat-anti rabbit IgG conjugated to Alexa Fluor 647 was used as secondary antibody. Imaging was performed using a confocal laser scanning microscope (Carl Zeiss, LSM710) with a 40× 1.2 NA objective (Carl Zeiss, 421767-9971-790).

Cranial window surgery and virus injections

For *in vivo* imaging experiments, we used 8–11-week-old male PCP×PBS hybrid mice that were heterozygous for PCP-GFP and homozygous for the Arc-PBS knock-in. Mice were anesthetized by intraperitoneal injection of ketamine/xylazine and fixed on a stereotactic frame. Chronic hippocampal windows were implanted as described previously (*3*). Briefly, a 2.7–2.8-mm diameter craniotomy was made at AP -2.0 mm, ML +1.8 mm from the bregma by using a 2.7-mm diameter trephine drill (FST) overlying the dorsal hippocampus. The dura was removed with forceps, and the cortical tissue above the CA1 region was carefully removed by aspiration. A customized stainless-steel cylindrical cannula with a 2.5-mm Ø round glass coverslip (Marienfeld, custom order) attached at the bottom was inserted and cemented to the skull using Meta-Bond (Parkell). Chronic cranial windows over the retrosplenial cortex (RSC) were implanted at AP -1.6 mm, ML +0.5 mm from the bregma. A 2.5-mm Ø round glass coverslip, was placed on the craniotomy and sealed with Meta-Bond. Before the Meta-Bond was cured, a customized stainless-steel head ring was placed around either the cannula or the window, and fixed by adding more Meta-Bond.

For virtual reality (VR) experiments, we injected 1 μl of $10^{13}$ vg/ml of AAV1-hSyn-NES-jRGECO1a (Addgene, #100854) into AP -2.0 mm, ML +1.8 mm and DV -1.4 mm from the bregma. Three days after virus injection, hippocampal window surgery was performed. The injection was made using a 33-gauge needle connected to a Hamilton syringe and mounted on a micro-injection pump (Harvard Apparatus, Pump 11 Pico Plus Elite).

Contextual fear conditioning (CFC) experiments

After cranial window surgery, mice were housed in the home cage for 7 days and habituated by daily handling and exposure to the anesthesia induction chamber for 5 min. On the eighth day (day 1 of the experiment), the mice were removed from the home cage and imaged under anesthesia (~1% isoflurane). On day 2, CFC was performed in Context A, which was a 250 mm × 250 mm × 250 mm acrylic box with white and black walls with yellow stripes. The chamber was equipped with a stainless-steel grid floor, an LED light, and a video camera (Canon, EOS Hi). The mice explored Context A for 180 s, and three 0.75 mA foot shocks of 2 s duration were delivered at 30 s intervals. From day 3, mice were returned to the conditioning chamber (Context A) or placed in a different context (Context B) for 180 s to assess freezing behavior induced by fear memory recall. Context B was an equilateral triangular chamber (360 mm per side) scented with 1% acetic acid. Contexts were cleaned with 70% ethanol before each session.

To assess freezing rates, we followed a method described previously (*4*), and wrote a custom MATLAB script that segments the mouse body and calculates the area of non-overlapping regions between each pair of consecutive images. If the non-overlapping area was below a threshold value for at least 0.5 seconds, the behavior was considered to be 'freezing.' The threshold value was adjusted until the freezing rate matched the manually obtained value.

Simulations

To compare the accuracy of GERI and conventional IEG promoter-based reporter expression methods for identifying IEG-positive neurons, we generated random *Arc* activation traces for 500 neurons. At the time of stimulation, 25% of the neurons were activated during 4 min to mimic the CFC experiment. The basal level of IEG activation was varied from 0–0.5% per min. The IEG activation traces were then convolved with the response function of GERI or that of shGFP reporter (*5*). The response function of GERI was obtained from the average intensity profile of the *Arc* transcription sites (n = 42) over time. The response function of shGFP was constructed from data in a previous report (*6*). The fraction of IEG-positive neurons was calculated by counting the number of neurons in which the intensity exceeded a threshold of

30% of the peak intensity at 6 min after stimulation for GERI and at 1 hr after stimulation for shGFP, respectively. The simulation was repeated 10 times. The accuracy was calculated by dividing the number of correctly predicted neurons by the number of total neurons (n = 500).

Electrical stimulation of cultured hippocampal neurons

Electrical burst stimulation was applied to cultured hippocampal neurons through two thin platinum wires. Hippocampal neurons were cultured from postnatal day 1 (P1) pups of WT mice, which were infected with AAV-hSyn-jRGECO1a at 14 days in vitro (DIV), and used for experiments at 17 DIV. Stimulation patterns were generated by an isolated pulse stimulator (A-M Systems, model 2100) and Arduino UNO microcontroller. Each burst consisted of two biphasic pulses with 2 ms duration, 10 ms interval, and 3.7 V amplitude. Ten bursts were applied at 6, 8, and 10 Hz with an interval of 7 s. A wide-field fluorescence microscope (Olympus, IX83) equipped with a 20× 0.5 NA objective (Olympus, UPLFLN20X), a motorized stage (Marzhauser), and an EMCCD camera (Andor iXon Life 888) was used to acquire time-lapse images of jRGECO1a at 30 frame per second (fps) during the electrical stimulation.

VR experiments

Water restriction and training were performed before VR experiments. Water restriction (1 ml/day) was applied for 5 days using a water dispenser connected to a multi-channel syringe pump (New Era, NE-1600). VR experiments were performed using the JetBall-TFT system (Phenosys), which consists of a 270° 6-panel monitor, a spherical treadmill, and a water reward device. To restrict the spherical treadmill movement to one dimension, a needle was inserted on one side of the treadmill. After 5 days of water restriction, training was performed on an infinite virtual linear track for 0.5~1 hour per day for 10-14 days, and water rewards were delivered at random positions. Mice were subjected to experimentation on the day following their demonstrated capability to travel longer than 50 m in 30 min. On experimental days 1 and 2, mice were exposed to virtual Context A (Fig. 4B) which was 3 m long, 0.4 m wide, and consisted of buildings and moon objects. On day 3, mice were placed into virtual Context B which was 3 m long, 1 m wide, and made up of objects and wall patterns different from Context A. After reaching the end of the track, the mice were teleported to the starting point. To induce a perception of movement on a continuous track, tunnels were placed at the front and the end of the virtual contexts. Water reward was delivered at the end of each tunnel.

*In vivo* two-photon imaging

*In vivo* imaging was performed using a two-photon excitation laser scanning microscope (Olympus, FVMPE-RS) equipped with two GaAsP photomultiplier tubes (PMTs), a Ti:Sapphire laser (Mai-Tai DeepSee, Spectra-Physics), a galvo/resonant scanner, and a 25× 0.95 NA water immersion objective with an 8-mm working distance (Olympus, XLSLPLN25XSVMP2). Excitation wavelengths of 900 nm (for *Arc* transcription imaging) and 1030 nm (for jRGECO1a imaging) were used, and fluorescence was collected via two PMTs after passing through a filter cube (Olympus, FV30-FGR) that consists of a 570 nm low-pass dichroic mirror and two emission filters (495–540 nm band-pass and 575–645 nm band-pass filters). For *Arc* transcription imaging after CFC, mice were anesthetized immediately after each behavior session by 5% isoflurane inhalation using a low-flow vaporizer (Kent Scientific, SomnoSuite) and mounted on the two-photon microscope. Anesthesia was maintained during imaging with 1~1.5% isoflurane, and body temperature was maintained at 37 °C. We scanned a volume of 250 μm × 250 μm × 20 μm at 1024 × 1024 × 81 voxels with a scan speed of 2 μs per pixel using a galvo scanner. To facilitate future relocation of regions of interest (ROIs), we marked 2~4 spots by laser ablation.

For *Arc* transcription and calcium imaging in virtual reality experiments, mice were mounted on the microscope while awake. *Arc* transcription imaging was performed for ~3 min before and after the VR experiment. We imaged 2 ROIs with a volume of 128 µm × 128 µm × 20 µm using a resonance scanner. During the virtual reality experiment, calcium activity was observed via jRGECO1a signal. Images of 256 µm × 256 µm areas were acquired at 30 Hz using a resonance scanner. Calcium imaging was initiated by triggering a signal from JetBall software to simultaneously record VR positions and calcium activity. After the VR experiment, we immediately anesthetized the mice to prevent additional stimulation, and re-imaged the ROI.

Fixed brain imaging

Fixed brain slices were prepared as previously described, with minor modifications (*7*). Mice were deeply anesthetized with isoflurane and perfused transcardially with 10-15 ml of PBS containing 10 U/ml heparin (Sigma, H3393), and 30-50 ml of fresh 4% paraformaldehyde (PFA, Sigma-Aldrich, 158127) in 0.1 M phosphate buffer (PB). Brains were post-fixed in 4% PFA at 4 °C overnight, and sectioned coronally to 50 µm with a vibratome (Leica, VT1200S). The sections were counterstained with 0.1 µg/ml DAPI (Invitrogen, D1306) in PBS and cover-slipped with VectaShield mounting media (Vector Labs, H-1400). Imaging was performed using a slide scanner (Zeiss, Axio Scan.Z1) with a 20× 0.8 NA objective (Zeiss, 420650-9902-000) or a confocal microscope (Zeiss, LSM780) with a 40× 1.3 NA oil immersion objective (Zeiss, 420460-9900-000).

To image jRGECO1a in fixed brain tissues, we used a wide-field fluorescence microscope (Olympus, IX83) equipped with a 20× 0.5 NA objective (Olympus, UPLFLN20X), a motorized stage (Marzhauser), and an EMCCD camera (Andor iXon Life 888). To obtain a large field of view, we performed 15 × 18 grid imaging and stitched the images after shading correction using BaSiC software (*8*).

Single-molecule fluorescence *in situ* hybridization (smFISH)

smFISH was performed using an RNAscope fluorescent multiplex assay (ACDBio) according to the manufacturer's protocol. After CFC, brains were harvested after decapitation and immediately frozen in -80 °C ethanol. The brains were sectioned coronally to 20 μm using a cryostat (Thermo Fisher Scientific, HM525 or Leica, CM1860) and collected on Superfrost microscope slides (Thermo Fisher Scientific, J1800AMNZ). The sections were fixed in 4% PFA at 4 °C for 15 min. Following serial dehydration in 50%, 70%, and 100% ethanol at RT, the sections were incubated in 100% ethanol overnight at -20 °C. Sections were then treated with proteinase IV for 30 min at RT and rinsed with 0.1 M PBS. Probes were applied for 2 h at 40 °C. Sections were incubated with probes for 2 h at 40 °C and subsequently incubated with amplifiers 1–4 at 40 °C. After counterstaining with DAPI solution, the sections were cover-slipped with ProLong mounting media (Invitrogen, P36965). Images were obtained using a Zeiss slide scanner with a 20× 0.8 NA objective or a confocal microscope (Zeiss, LSM780) with a 40× 1.3 NA oil immersion objective (Zeiss, 420460-9900-000). Probes used in this study were Arc-C1 (Cat# 316911), PP7-C2 (Cat# 300031, custom-designed), c-Fos-C2 (Cat# 316921), and Egr-1-C3 (Cat# 423371).

Image analysis and calcium signal extraction

For *Arc* mRNA imaging after CFC, image registration and analysis were performed with custom-written MATLAB codes (Fig. S2). The MATLAB scripts used for image registration and analysis are available at GitHub (https://github.com/Neurobiophysics). Motion artifacts from breathing and cardiac function were corrected by using an auto-fluorescence image (red channel) as a reference. Images taken on different days were aligned by rotating and translating the images. After the alignment, we subtracted the red channel image from the green channel image after normalization by maximum values to remove auto-fluorescence signals. In our PP7-GFP system, the nuclear localization sequence was added to PCP-GFP to facilitate the identification of neuronal nuclei by the GFP signal. The 3-dimensional (3D) coordinates of the cell centroids were determined by using a circle-finding algorithm (*9*) in the XY and XZ plane. By examining the z-section of each cell image, two experimenters blindly classified the neurons into three groups: neurons with *Arc* transcription sites (*Arc*+), neurons without *Arc* transcription sites (*Arc*-), and not-determined (ND) cells. Neurons that showed one or two bright spots in the same XY position in at least two consecutive z-slices were classified as *Arc*+ cells. ND cells include cropped cells and cells that had PCP-GFP expression levels that were either excessive or inadequate to detect transcription sites. Any disagreement between the two experimenters was resolved by a third person.

For the analysis of immunohistochemistry, the 3D coordinates of the cell centroids were determined by using a circle-finding algorithm (*9*). Arc protein-positive and negative neurons were manually classified by the brightness of Arc protein signal. Then the Arc protein intensity threshold was determined by fitting the intensity histograms of Arc-positive and negative cells with normal distributions. Then all brain slices were analyzed based on the same threshold.

Particles in dual-color smFISH images were detected using FISH-quant software (*10*). FISH-quant provided sub-pixel positions and intensity by performing three-dimensional Gaussian fitting. A custom-written MATLAB script classified particles into single mRNA and transcription sites by setting thresholds for amplitude and width of three-dimensional Gaussian function (Fig. S3C). Threshold values were adjusted by examining each slice image manually. Transcription sites within 0.3 μm detected in the Atto 550 dye (CDS target) and Atto 647 dye (PBS target) channels were considered identical (Fig. S3D). To quantify the number of nascent mRNAs in a transcription site, we calculated the intensity of particles by $I = Amplitude * \sigma_x * \sigma_y * \sigma_z$ which is an integrated value of a three-dimensional Gaussian function, and divided the intensity by the median intensity of single mRNAs.

The image registration and data extraction process for the VR experiment included: 1) motion correction of the *Arc* mRNA images using NoRMCorre (*11*) and stackReg (*12*) software; 2) stitching two ROIs and subtracting auto-fluorescence; 3) alignment of images taken before and after VR and on different days; 4) matching the same field of view in *Arc* mRNA and calcium images; 5) obtaining nucleus centroid coordinates using Imaris software (Bitplane); 6) calcium source extraction using CaImAn software(*13*), which is based on a constrained non-negative matrix factorization (CNMF) algorithm by seeding the nucleus centroid locations as initial spatial components; and 7) matching the nucleus and spatial components detected by CaImAn finding the nearest centroid position.

To detect transcription sites that are near diffraction-limited spots, we needed a high signal-to-noise ratio, high spatial resolution, and little distortion from motion artifacts. For this, we acquired 45 images in each z-plane at 30 Hz and obtained the average image after image registration by x-y translation using NoRMCorre. The z-axis of the image was then aligned using StackReg software. Two ROIs were stitched using the stitching plugin (*14*) in ImageJ software. To align stitched voxels taken in different days, we wrote a custom MATLAB script that finds overlapping voxels by calculating cross-correlations (Fig. S7). The calcium activity images were first corrected for motion artifacts by using NoRMCorre software. Cross-correlations were then calculated between the *Arc* mRNA images and averaged calcium images to find the same field of view and corresponding z-plane. To obtain the 3D coordinate of neuronal nuclei, we used the surface segmentation tool of Imaris (Bitplane) software. The locations of cells within 10 μm of the calcium imaging plane were used as the initial spatial components in CaImAn software. After running CaImAn, the output spatial components were used to identify nuclear coordinates by finding the nearest neighbor centroid position. The calcium activity traces were calculated using the temporal component values from CaImAn, and relative changes (ΔF/F) were computed. The calcium trace baseline was selected manually and corrected for photobleaching. A calcium transient event was defined as an event starting from the moment ΔF/F exceeded three standard deviations (SD) of the baseline to the moment that ΔF/F returned to within 0.5 SD of the baseline. We considered cells with at least one calcium transient to be active cells. False-positive calcium transients due to motion artifacts in the z-axis were occasionally observed during murine grooming behavior. To remove these artifacts, we used the calcium activity data recorded only during walking (speed > 0.5 cm/s and run length > 0.5 cm). To resolve finer spiking activity, we deconvoluted ΔF/F traces and inferred the spike traces (*15*). $Ca^{2+}$ event rates were calculated by dividing the sum of the inferred spikes by time. A burst was defined as spikes that were larger than a threshold value of 2.5 in a 33-ms time bin or continuous over multiple time bins. We defined theta-burst events as bursts occurred at interburst intervals of 100 to 167 ms (6 to 10 Hz) following a similar method in (*16*).

Place cell identification

Place cells were defined using a method similar to an approach described previously (*3*). The 3-m-long virtual track was first divided into 150 position bins (2 cm per bin). Using the ΔF/F traces obtained during running (speed >1.5 cm/s and run length > 5 cm), we calculated the sum of ΔF/F for each bin and normalized by the dwell time in each bin to generate a ΔF/F field map. The resulting ΔF/F field map was smoothed by computing the moving average with a sliding window of three bins. The potential place field was identified as a region exceeding the median value of the ΔF/F field map. Place cells were then identified using the following criteria: 1) the potential field is wider than 20 cm; 2) the mean value of the ΔF/F field map in the potential field must be 2.2 times higher than the mean value of the ΔF/F field map outside the potential field; and 3) $Ca^{2+}$ events must be present in at least 15% of the visits to the potential field. To evaluate whether each mouse actually distinguished virtual Contexts A and B, we calculated spatial correlation. For neurons identified as place cells on day 1, we calculated the Pearson's correlation coefficient of the ΔF/F field map between the same or different contexts. The spatial information was calculated using the following formula (*17*):

$$\text{Spatial information} = \sum_i \frac{\lambda_i}{\lambda} \log_2 \frac{\lambda_i}{\lambda} P_i$$

where $\lambda$ is the $Ca^{2+}$ event rate, $\lambda_i$ is the mean $Ca^{2+}$ event rate in the $i^{th}$ place bin, and $P_i$ is the probability that the mouse stays in the $i^{th}$ place bin.

Network graph

We first converted the $Ca^{2+}$ event traces into binary and temporally binned (1 sec per bin) traces. The resulting traces were used to generate a correlation matrix by calculating Pearson's correlation coefficients. We then generated a binary adjacent matrix with pairs that were statistically significant ($p < 0.05$) and had a correlation coefficient higher than 0.1. The network graph was visualized using Gephi (https://gephi.org/) software with the ForceAtlas2 layout. We calculated degrees, cluster coefficients, and modularity using custom written MATLAB scripts. The degree $k_v$ denotes the number of edges in each neuron. The normalized degree was calculated by dividing $k_v$ by (the number of neurons in network -1). Clustering coefficients were calculated using the following formula (*18*):

$$C_v = \frac{\sum_{j,k} A_{vj} A_{jk} A_{kv}}{k_v(k_v - 1)}$$

where $A$ is the adjacent matrix in which $A_{ij}$ is a binary value indicating whether the i$^{th}$ neuron and the j$^{th}$ neuron are correlated. The normalized modularity was calculated using the following formula (*19*):

$$Q_{normalized} = \frac{Q}{Q_{max}} = \frac{\sum_{i,j}\left(A_{ij} - \frac{k_i k_j}{2m}\right)\delta(t_i, t_j)}{2m - \sum_{i,j}\frac{k_i k_j}{2m}\delta(t_i, t_j)}$$

where $m$ is the number of edges in the network, $t_i$ denotes the module (*Arc*++/non-*Arc*++ or place cell/non-place cell) of the i$^{th}$ neuron, and $\delta$ is a delta function.

### Supplementary Figures

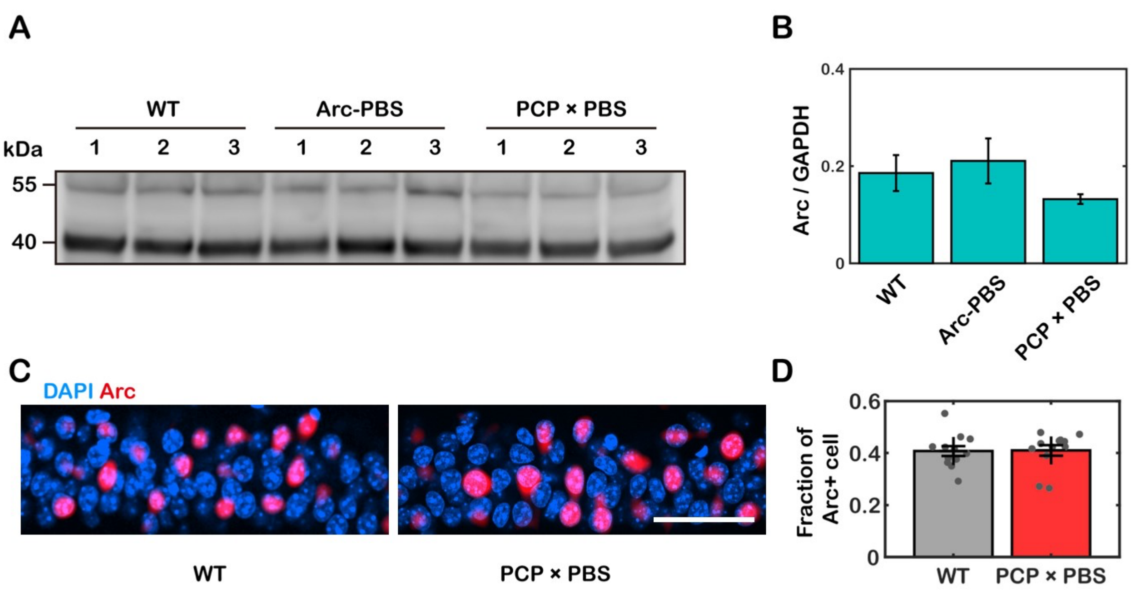


**Fig. S1. Comparison of Arc protein expression levels. (A)** Western blot of Arc (55 kDa) and GAPDH protein (36 kDa) in brain tissue lysates of three mice from each group of wild type (WT), homozygous Arc-PBS knock-in (Arc-PBS), and double homozygous Arc-PBS knock-in × PCP-GFP (PCP×PBS). **(B)** Quantification of relative Arc protein expression levels using GAPDH as a loading control. No significant differences were observed between the mouse lines (n = 3 mice for each line). Error bars represent standard deviation (SD). **(C)** Example images of Arc immunofluorescence (red) in dorsal CA1 of WT (left) and PCP×PBS mice (right). Scale bar, 50 µm. **(D)** The fraction of Arc-expressing cells in WT and PCP×PBS mice (n = 12 slices from two mice for each mouse line). Error bars represent the standard error of the mean (SEM).

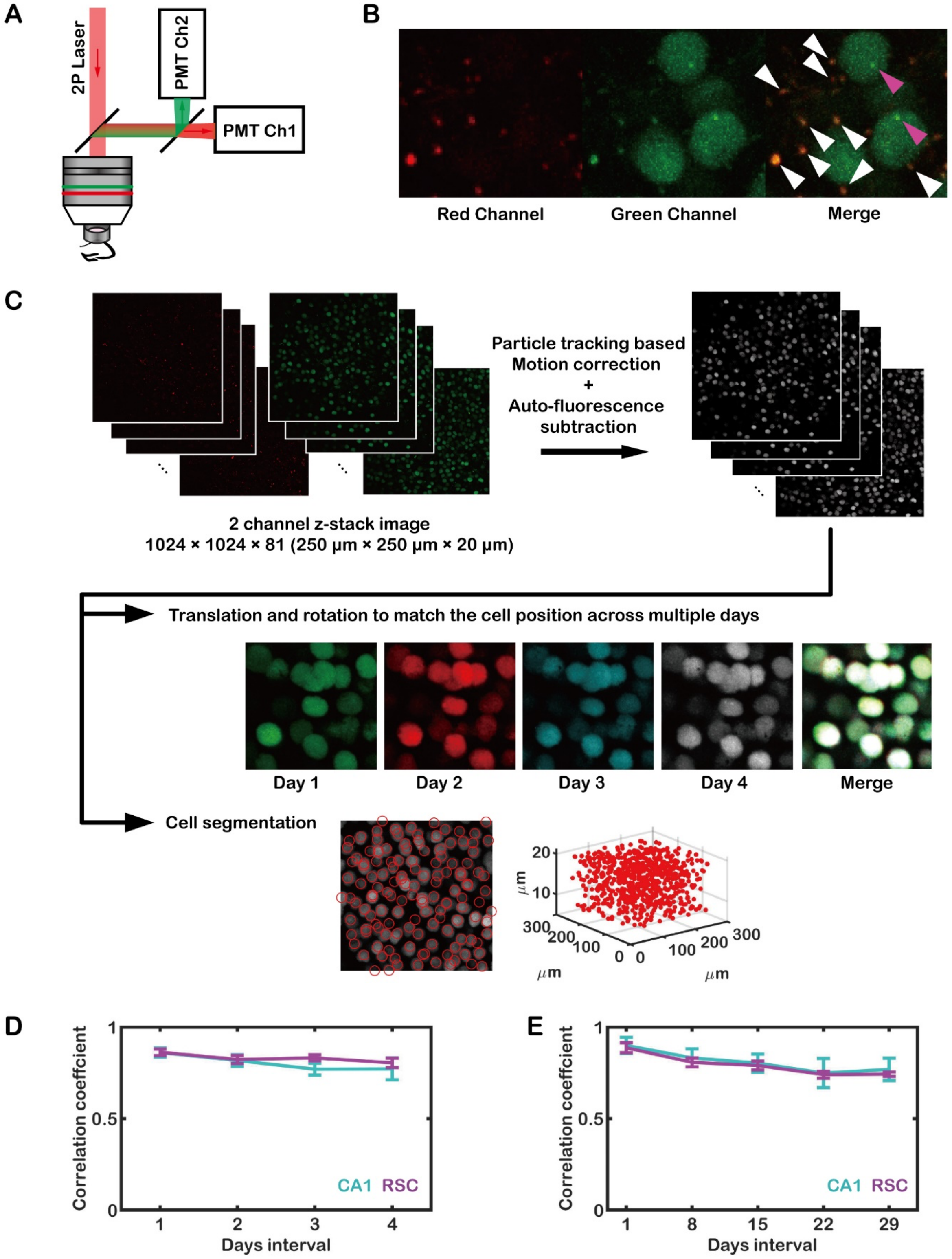


**Fig. S2. Image processing procedures. (A)** Two-photon microscopy for dual-color *in vivo* imaging. **(B)** Raw images taken by two-photon microscopy. Auto-fluorescent particles (white arrow) were observed in both green and red channels. *Arc* transcription sites (magenta arrow) were detected only in the green channel. **(C)** Motion artifacts were corrected by a custom particle-tracking algorithm. Auto-fluorescent signals were subtracted and the images were registered across multiple imaging sessions by translation and rotation. Finally, automatic cell segmentation was performed to find neuronal coordinates. **(D-E)** Pearson's correlation coefficients were calculated between images taken 1–4 days apart (D) and 1–4 weeks apart (E). Error bars represent the SEM.

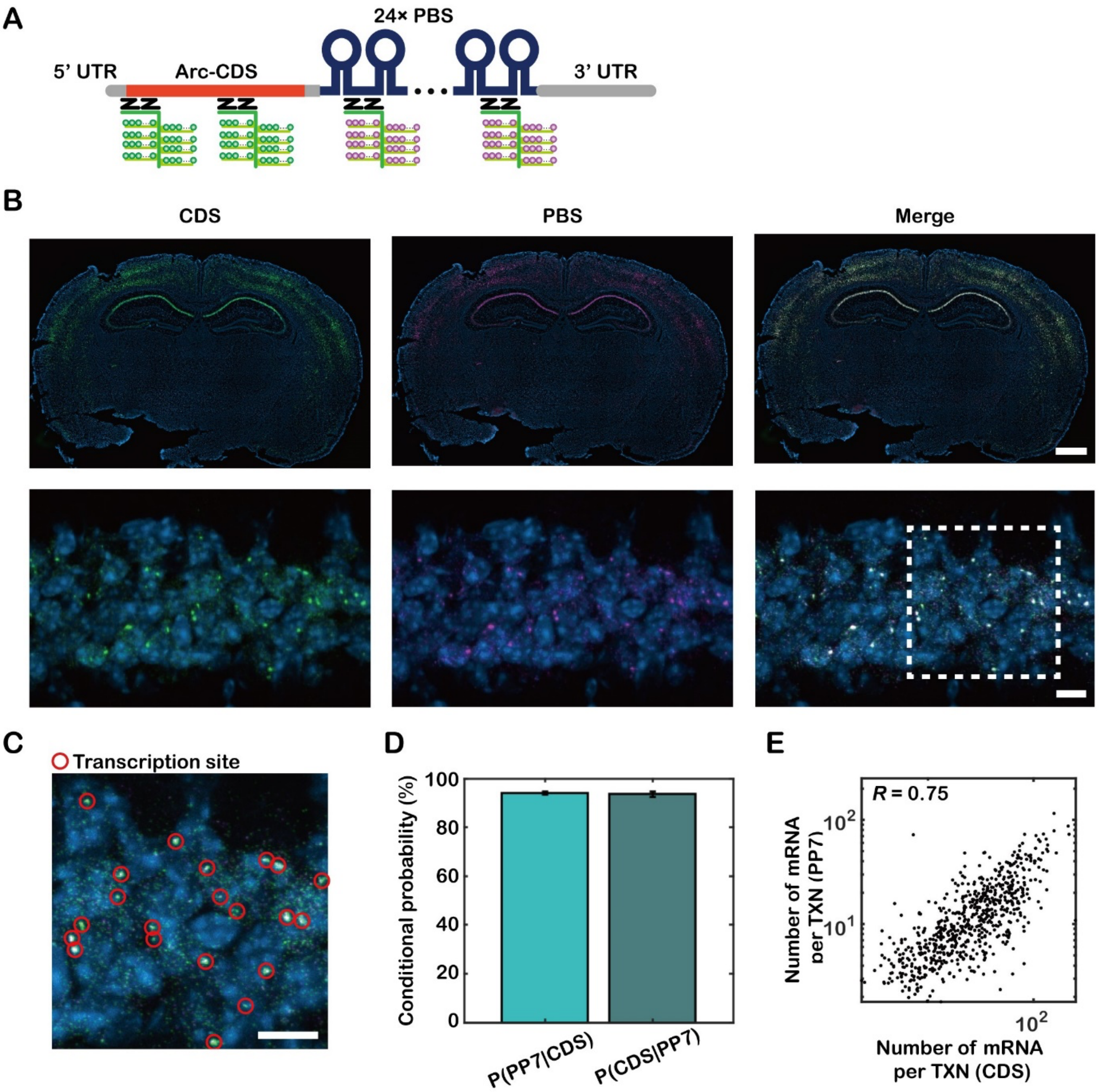


**Fig. S3. Dual-color single-molecule fluorescence *in situ* hybridization (smFISH) targeting the CDS and PBS region of *Arc* mRNA. (A)** Schematic for dual-color smFISH. The CDS region of *Arc* mRNA was detected by Atto 550 dye (green) and PBS was detected by Atto 647 dye (magenta). **(B)** Representative smFISH images. **(C)** An enlarged image of the dotted box in (B). Red circles denote transcription sites. **(D)** Conditional probability of detecting transcription sites in different channels. **(E)** Scatter plot of the number of nascent mRNAs per transcription site detected by the probes targeting CDS and PBS ($n$ = 670 transcription sites, correlation calculated by Spearman's coefficient $R$). Scale bars, (B, upper panels) 1 mm and (B, lower panels and C) 10 μm.

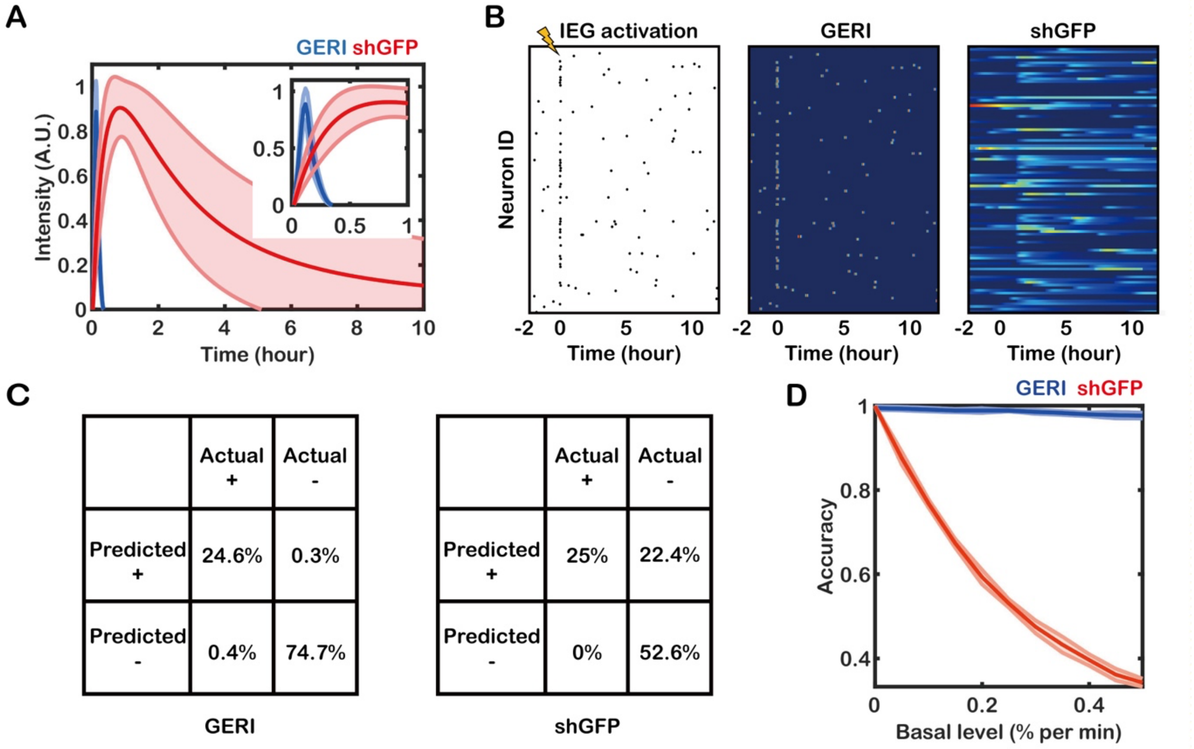


**Fig. S4.** Simulation of GERI and shGFP reporter-based detection of IEG+ neurons. (A) The response functions of GERI (blue) and shGFP (red) over time. The inset shows a magnified view of the first hour. (B) Example images of simulated IEG activation traces (left), and intensity traces generated by convolution of the activation trace and the response function of GERI (middle) and shGFP (right). The probability of IEG activation was 0.1% per min in the baseline and 25% in 4 min (6.25% per min) during stimulation (lightning symbol). (C) Confusion matrices showing the classification results using GERI (left) and shGFP (right) when the basal IEG activation probability was 0.1% per min and stimulated activation probability was 25% in 4 min. (D) The accuracy of GERI (blue) and shGFP (red) methods plotted for different basal levels of IEG activation.

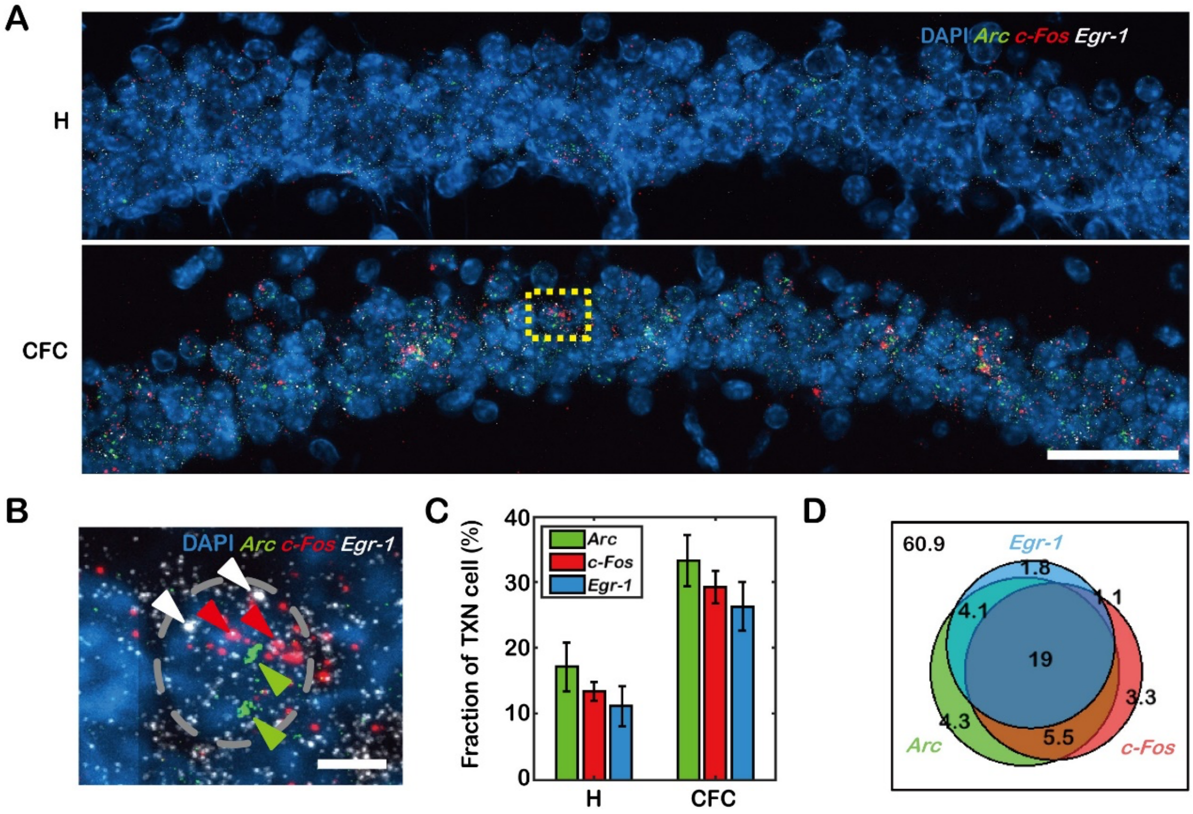


**Fig. S5. Three-color smFISH targeting *Arc, c-Fos* and *Egr-1* mRNA in CA1. (A)** Representative smFISH images of hippocampi from mice held in the home cage (H; upper panel) and after CFC (lower panel). *Arc, c-Fos* and *Egr-1* mRNA were labeled with FITC (green), Atto 550 (red), and Atto 647 (white) dyes, respectively. **(B)** Enlarged image of yellow box in (A). White, red, and green arrows indicate transcription sites of *Egr-1*, *c-Fos*, and *Arc*, respectively. **(C)** The fraction of cells with a transcription site for each gene. **(D)** A Venn diagram showing the percentage of cells expressing each gene. Scale bars, (A) 50 µm and (B) 5 µm. Error bars represent the SEM.

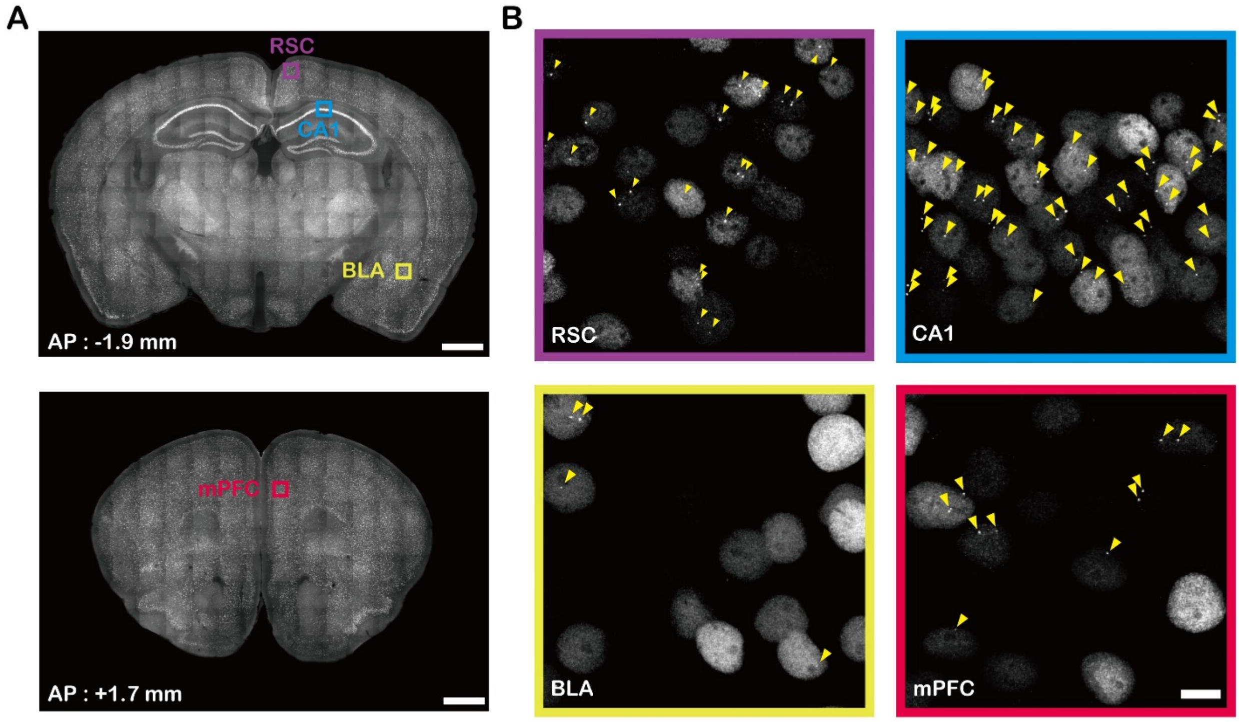


**Fig. S6. GFP images of fixed brain slices. (A)** Coronal sections of a PCP×PBS mouse brain at two AP positions (AP = -1.9 mm and +1.7 mm). Upper panel includes retrosplenial cortex (RSC), CA1, and basolateral amygdala (BLA). Lower panel includes medial prefrontal cortex (mPFC). **(B)** Enlarged images of the inset boxes in (A). Scale bars, (A) 1 mm and (B) 10 µm.

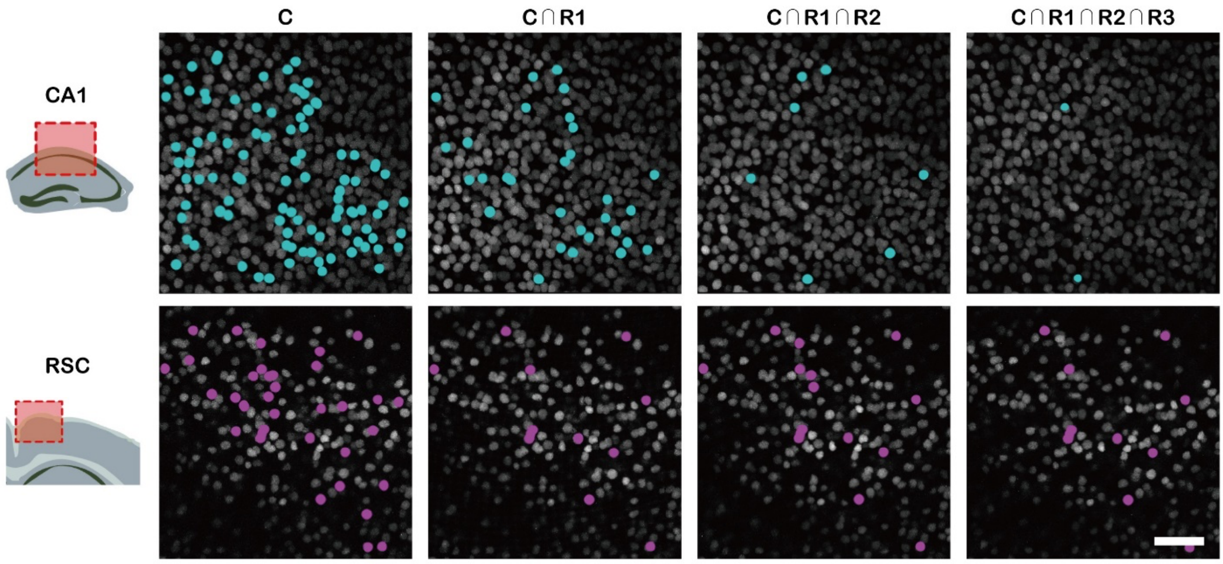


**Fig. S7. Overlap of *Arc*+ neurons upon recent memory retrieval**. Representative images of CA1 (upper panels) and the RSC (lower panels) showing overlapping populations of *Arc*+ neurons upon recent memory retrievals. (cyan dots, CA1; magenta dots, RSC). Scale bar, 50 μm.

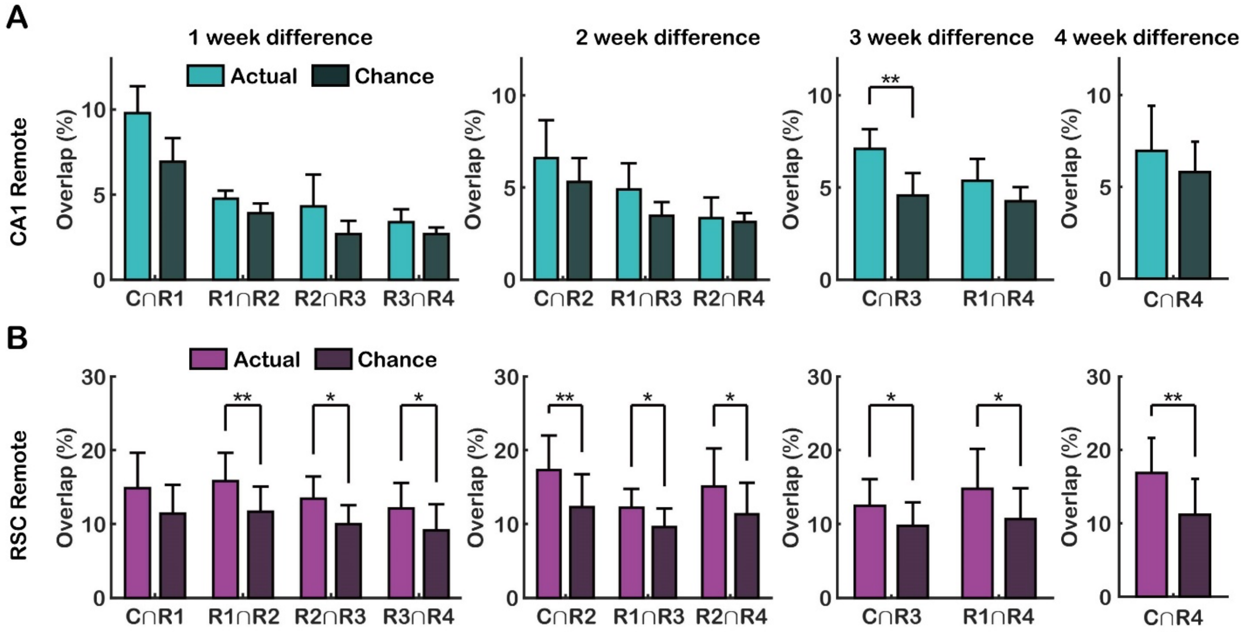


**Fig. S8. Overlap between *Arc*+ populations after remote memory retrievals (A-B)** The percent overlap between *Arc*+ ensembles in CA1 (A) and the RSC (B), with 1–4-week differences compared with the chance level (* $P < 0.05$, ** $P < 0.01$ by one-tailed pairwise $t$ test). RSC ensemble overlap was generally significantly greater than chance levels, whereas most CA1 ensembles exhibited chance-level overlap. Error bars represent the SEM.

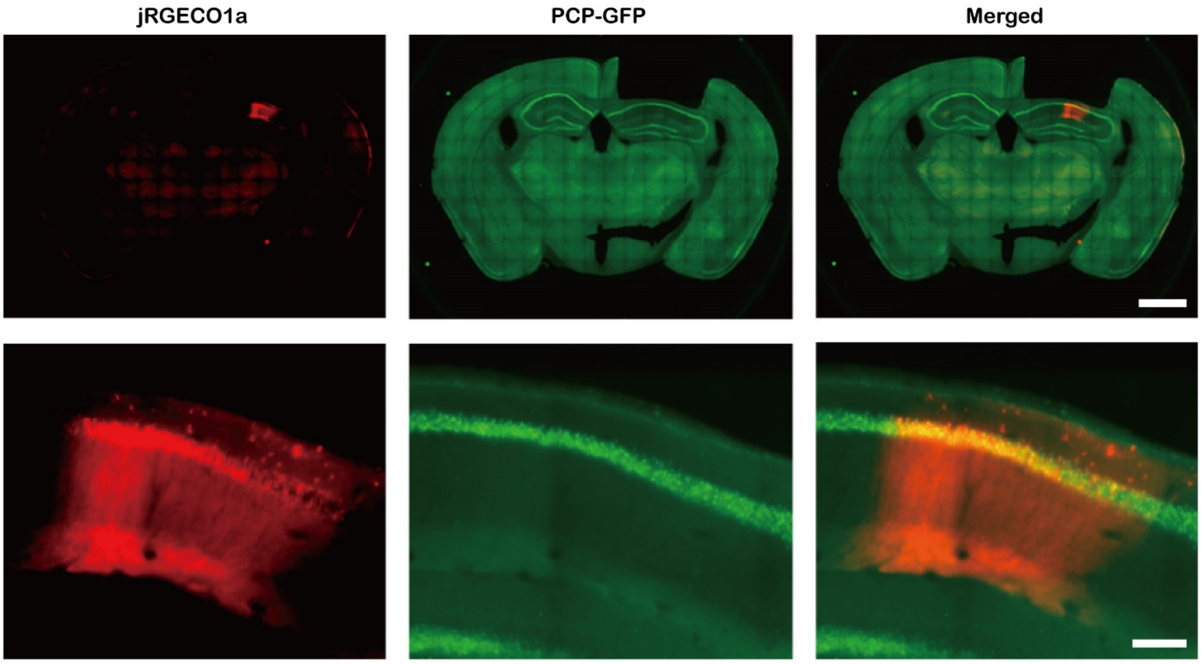


**Fig. S9. Expression of jRGECO1a in CA1 of PCP×PBS mouse.** AAV1-hSyn-NES-jRGECO1a was injected into the dorsal CA1 of PCP×PBS mice**.** Coronal section images of the dorsal CA1 pyramidal cell layer expressing jRGECO1a (red) and PCP-GFP (green). Scale bars, (upper panel) 1 mm and (lower panel) 100 µm.

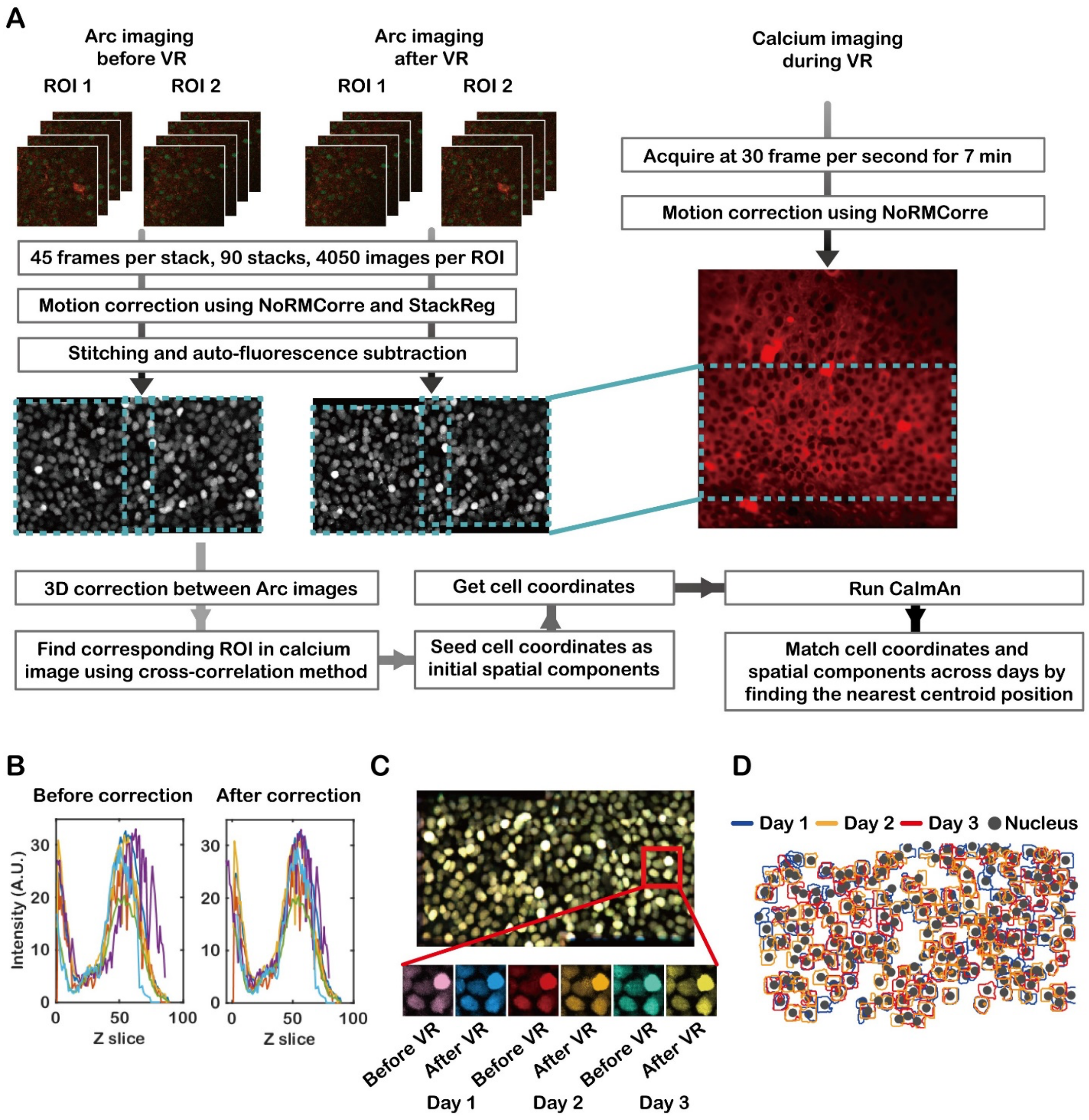


**Fig. S10. Image registration pipeline for *Arc* mRNA and calcium images. (A)** Raw images of *Arc* mRNA were first corrected for motion artifacts, followed by ROI stitching, auto-fluorescence subtraction, correction across days, and cell coordinate detection. Calcium images were corrected for motion artifacts, and calcium footprints were detected using constrained non-negative matrix factorization (CNMF) with information of detected cells (see Methods for details). **(B)** A z-slice profile of *Arc* mRNA images before (left) and after correction (right).

**(C)** Corrected images of *Arc* mRNA across multiple days. **(D)** Identified spatial footprints using CNMF over multiple days. Cell coordinates and spatial footprints were matched by finding the nearest centroid position.

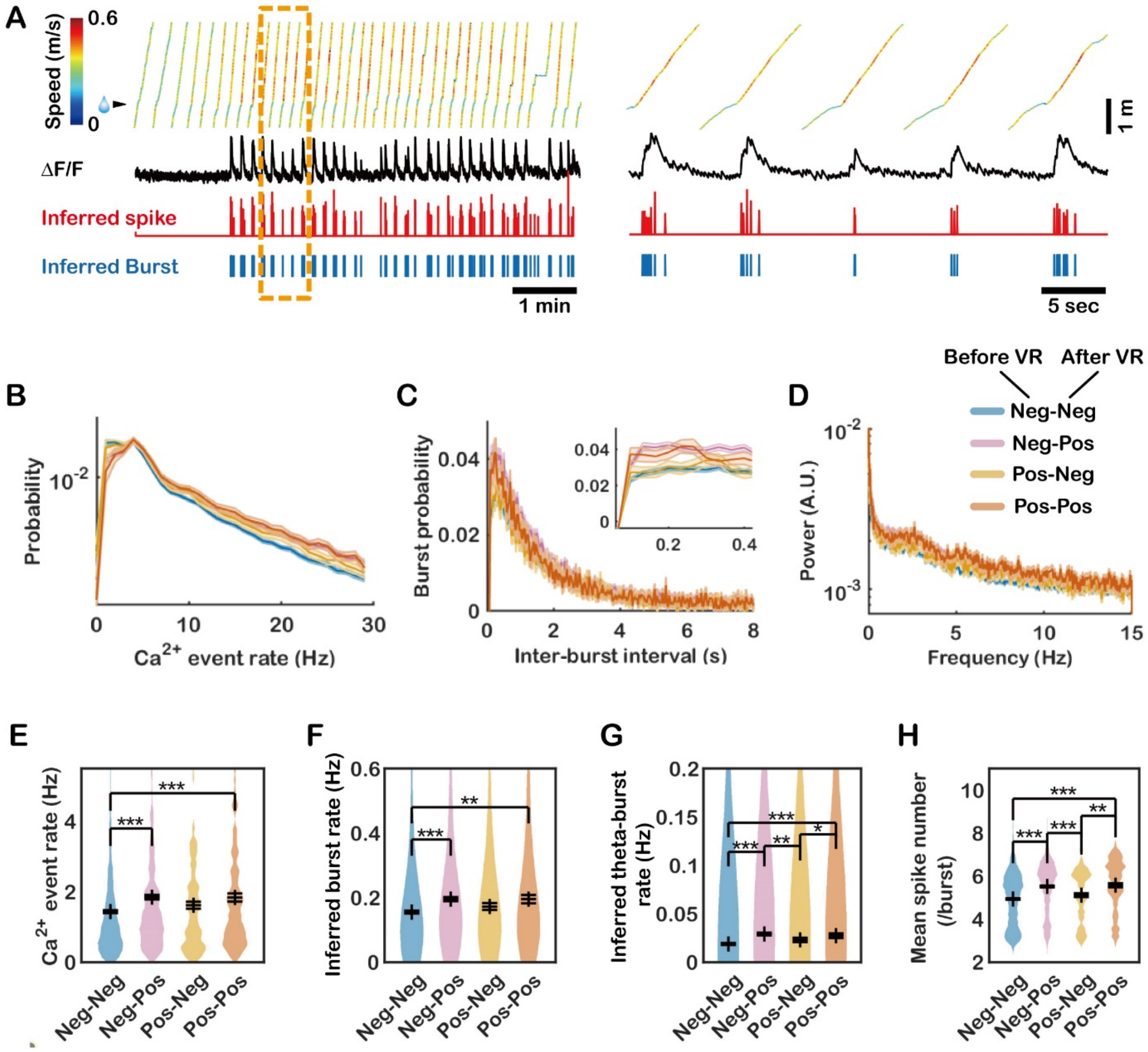


**Fig. S11. Calcium activity of CA1 neurons during VR navigation. (A)** An example of the VR track position with the colormap of the running speed over time. A representative neuron's calcium trace, inferred spike trace from deconvolution, and inferred burst trace are shown at the bottom (left). A segment of traces in the yellow dashed box is magnified at right. **(B-D)** We classified neurons into four classes according to neuronal transcription of *Arc* mRNA before and after VR. The $Ca^{2+}$ event rate histogram (B), autocorrelation (C), and power spectrum (D) of each class were plotted. **(E-H)** The $Ca^{2+}$ event rate, inferred burst rate, inferred theta-burst rate, and mean spike number per burst of each group were plotted (* $P < 0.05$, ** $P < 0.01$, ***

$P < 10^{-10}$ by rank-sum test). Generally, neurons in the Neg-Pos group had similar activity properties to neurons in the Pos-Pos group. Error bars represent the SEM.

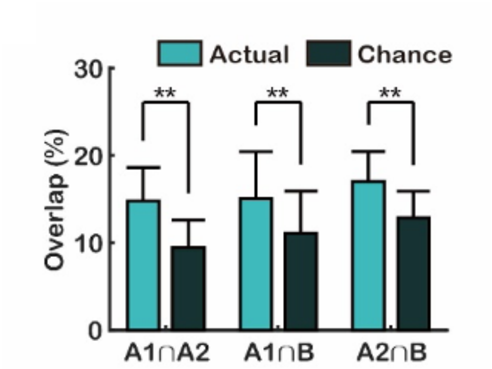


**Fig. S12. Comparison of overlap rates with chance levels.** Overlap between *Arc*+ neurons identified on days 1, 2, and 3 compared with chance levels (** $P < 0.01$, by pairwise $t$ test). Error bars represent the SEM.

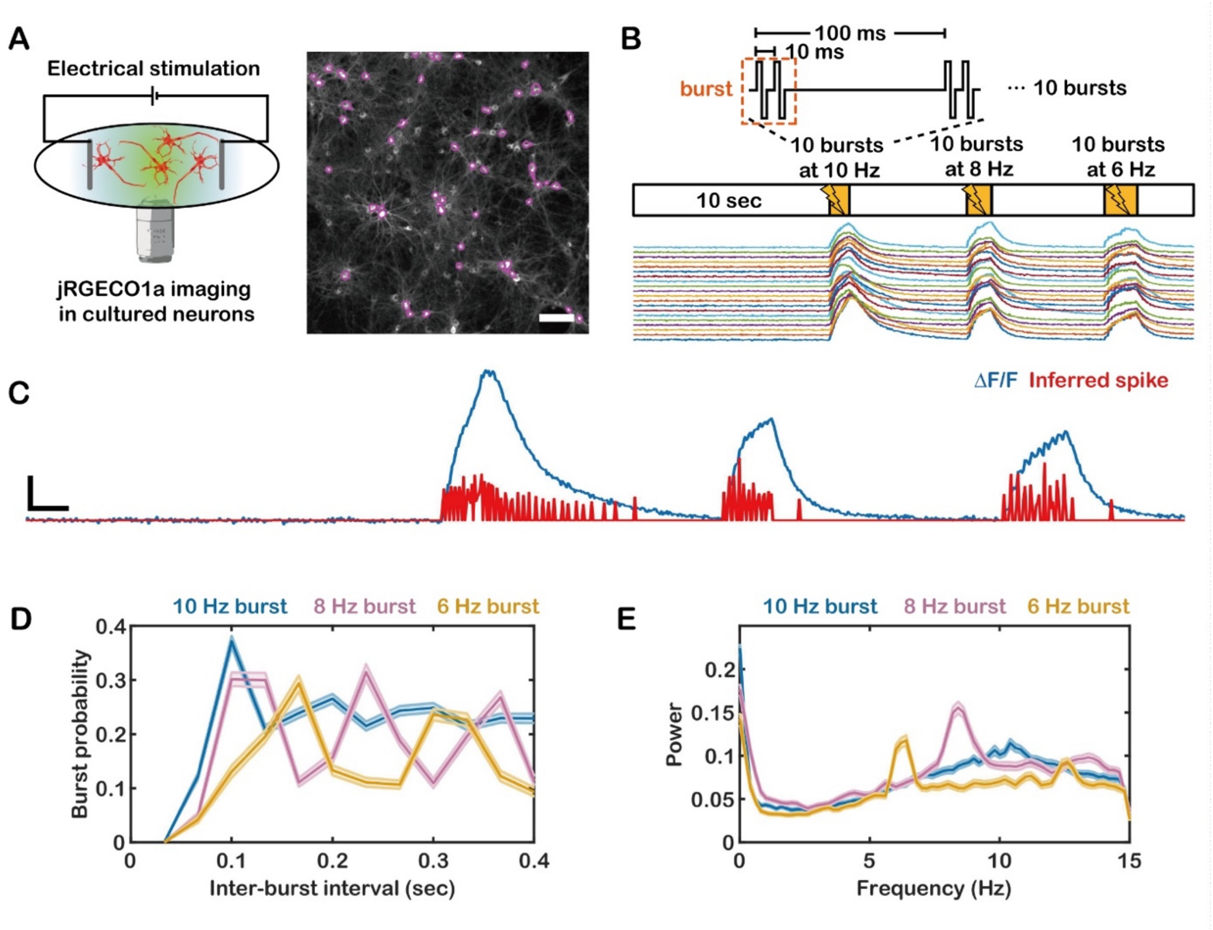


**Fig. S13. Calcium imaging of primary cultured hippocampal neurons upon 6–10 Hz burst electrical stimulations. (A)** Experimental scheme of electrical stimulation (left) and a representative image of cultured neurons expressing jRGECO1a (right). Magenta contours represent the ROIs for calcium fluorescence signals. **(B)** A single burst contains two electrical pulses with 10 ms interval. 10 bursts were delivered to neurons at 10, 8, and 6 Hz interspaced by 7 s. The calcium traces ΔF/F of 20 representative neurons are shown at the bottom. **(C)** An example ΔF/F trace (blue) and inferred spike trains (red). **(D)** Histogram of inter-burst intervals obtained from the inferred spike trains during 10, 8, and 6 Hz stimulation. **(E)** Power spectra of inferred bursts during 10, 8, and 6 Hz stimulation. Scale bars, (A) 100 µm, (C, vertical) 500% ΔF/F and (C, horizontal) 1 s.

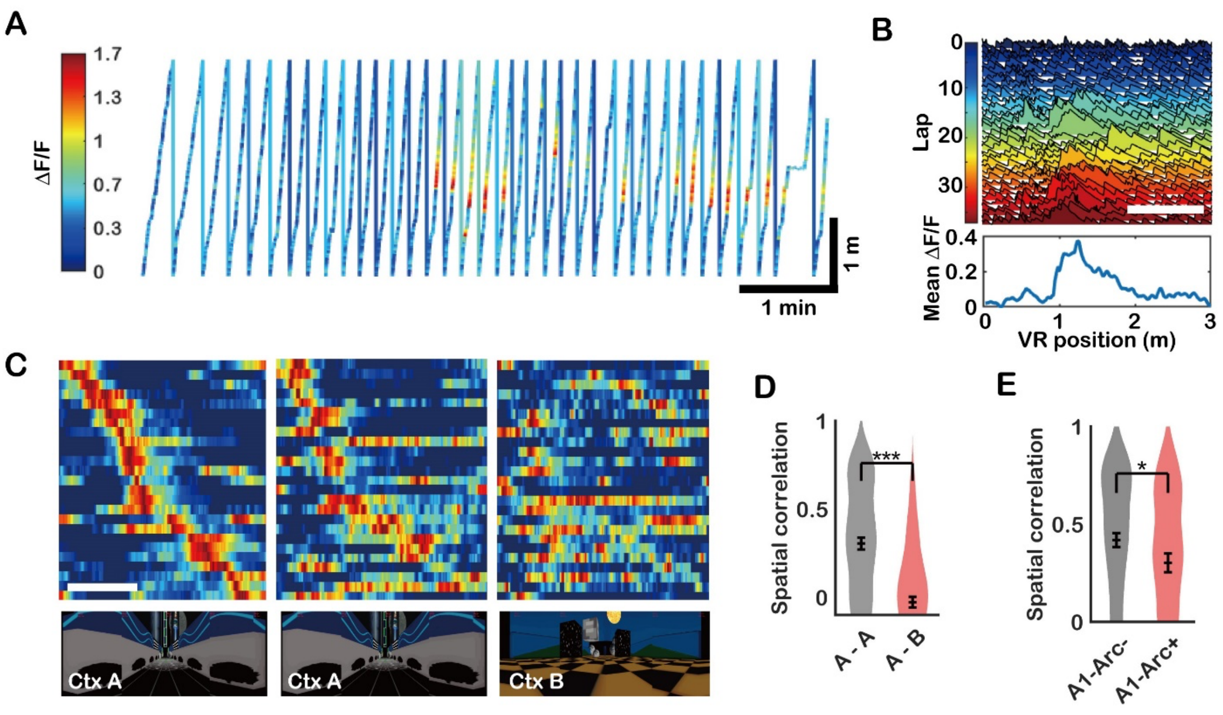


**Fig. S14. Spatial correlation after exposure to the same or different virtual contexts. (A)** Position versus time plot of a mouse running along the virtual linear track, colored according to ΔF/F values of a representative place cell. **(B)** Joyplot showing the calcium traces of each lap and the mean calcium trace of the place cell shown in (A). **(C)** Mean place fields on days 1–3, sorted by place cells detected on day 1. **(D)** Spatial correlation between place fields when mice were exposed to the same (A-A) or different (A-B) contexts. **(E)** Comparison of A1-*Arc*- and A1-*Arc*+ neurons in terms of their spatial correlation between place fields when mice were exposed to the same (A-A) contexts. Scale bar, (C) 1 m.

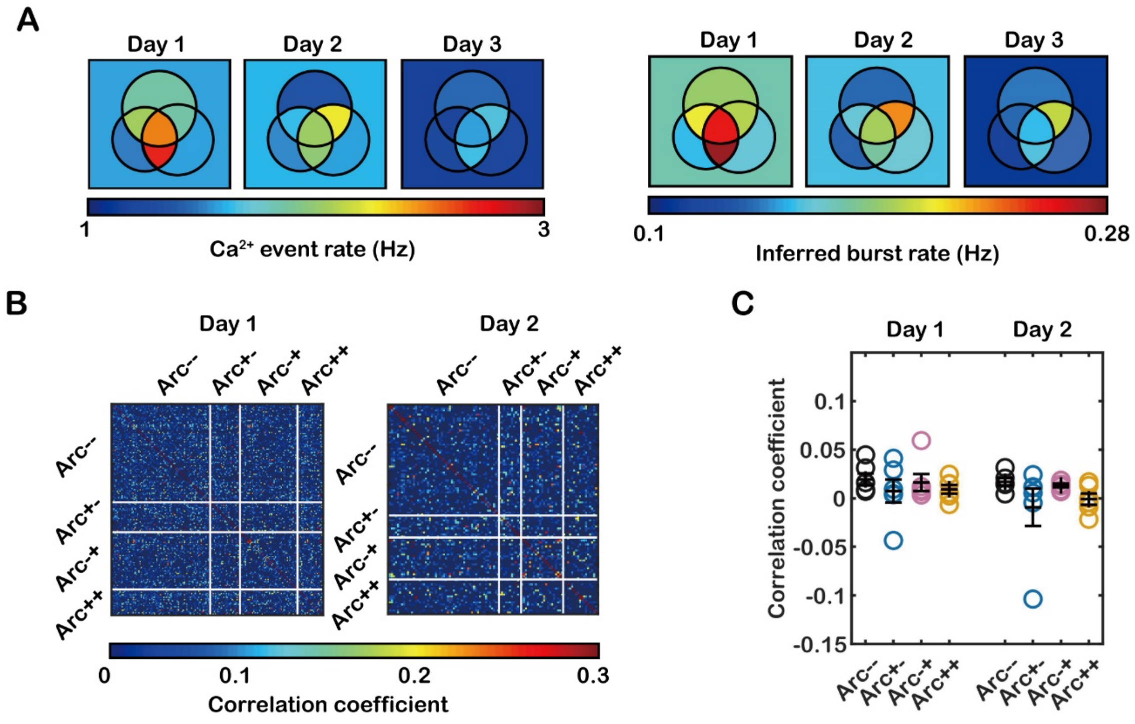


**Fig. S15. Ca$^{2+}$ event rates, inferred burst rates and correlation coefficients of *Arc*+ subpopulations. (A)** Venn diagrams of *Arc*+ neurons on days 1, 2, and 3 colored by the Ca$^{2+}$ event rate (left) and the inferred burst rate (right). **(B)** Correlation matrices generated by calculating the Pearson's correlation coefficient between binned spike traces. The neurons were arranged in the order of Arc--, Arc+-, Arc-+ and Arc++ neurons. **(C)** The correlation coefficient of the binned spike traces of the neurons in each group ($n$ = 6 mice).

### Supplementary Movie Captions

**Movie S1. Three-dimensional view of CA1 in a live PCP×PBS mouse.** A scanning volume of 250 μm × 250 μm × 20 μm was imaged in 1024 × 1024 × 81 voxels with a scan speed of 2 μs per pixel.

**Movie S2. Time-lapse calcium images of CA1 neurons during VR navigation.** The calcium activity of neurons in an area of 256 μm × 256 μm was recorded at 30 fps. The orange contours indicate the *Arc*++ neurons. The movie is played at 16 times real speed.
